# Human aneuploid cells depend on the RAF/MEK/ERK pathway for overcoming increased DNA damage

**DOI:** 10.1101/2023.01.27.525822

**Authors:** Johanna Zerbib, Marica Rosaria Ippolito, Yonatan Eliezer, Giuseppina De Feudis, Eli Reuveni, Anouk Savir Kadmon, Sara Martin, Sonia Viganò, Gil Leor, James Berstler, Kathrin Laue, Yael Cohen-Sharir, Simone Scorzoni, Francisca Vazquez, Uri Ben-David, Stefano Santaguida

**Affiliations:** Department of Human Molecular Genetics and Biochemistry, Faculty of Medicine, Tel Aviv University, Tel Aviv, Israel; Department of Experimental Oncology at IEO, European Institute of Oncology IRCCS, Milan, Italy; Broad Institute of MIT and Harvard, Cambridge, MA, USA; Department of Oncology and Hemato-Oncology, University of Milan, Milan, Italy

## Abstract

Aneuploidy is a hallmark of human cancer, yet the cellular mechanisms that allow cells to cope with aneuploidy-induced cellular stresses remain largely unknown. Such coping mechanisms may present cellular vulnerabilities that can be harnessed for targeting cancer cells. Here, we induced aneuploidy in non-transformed RPE1-hTERT cells and derived multiple stable clones with various degrees of chromosome imbalances. We performed an unbiased genomic profiling of 6 isogenic clones, using whole-exome and RNA sequencing. We then functionally interrogated their cellular dependency landscapes, using genome-wide CRISPR/Cas9 screens and large-scale drug screens. We found that aneuploid clones activated the DNA damage response (DDR), and were consequently more resistant to further DNA damage induction. Interestingly, aneuploid cells also exhibited elevated RAF/MEK/ERK pathway activity, and were more sensitive to several clinically-relevant drugs targeting this pathway, and in particular to genetic and chemical CRAF inhibition. CRAF activity was functionally linked to the resistance to DNA damage induction, as CRAF inhibition sensitized aneuploid cells to DNA damage-inducing chemotherapies. The association between aneuploidy, RAF/MEK/ERK signaling, and DDR was independent of p53. The increased activity and dependency of aneuploid cells on the RAF/MEK/ERK pathway was validated in another isogenic aneuploid system, and across hundreds of human cancer cell lines, confirming their relevance to human cancer. Overall, our study provides a comprehensive resource for genetically-matched karyotypically-stable cells of various aneuploidy states, and reveals a novel therapeutically-relevant cellular dependency of aneuploid cells.

## Introduction

Aneuploidy, an imbalanced number of chromosomes, is a unique characteristic of cancer cells^1–3^. In order to selectively target aneuploid cancer cells, it is imperative to better understand the molecular, cellular and physiological consequences of this phenomenon. Whereas many of the effects of aneuploidy are chromosome-specific, the aneuploid state itself is associated with cellular stresses that aneuploid cells must overcome in order to survive and proliferate^4,5^. Uncovering the cellular coping mechanisms of aneuploid cells could enable their selective targeting.

So far, attempts to study aneuploidy in human cells have mostly focused on non-isogenic tumors^6^ and cell lines^7^. For example, we have recently mapped the aneuploidy landscapes of ∼1,000 human cancer cell lines and revealed an increased vulnerability of aneuploid cancer cells to inhibition of the spindle assembly checkpoint and of the mitotic kinesin *KIF18A*^7^. However, such comparisons may be inherently confounded by aneuploidy-associated genomic features and other differences between non-isogenic cancer samples. Attempts to generate matched (pseudo-)diploid and aneuploid cell models have also been reported, mostly on p53-mutant and chromosomally unstable genetic backgrounds^8,9^. Karyotypically stable p53-WT models have been generated as well, but these models used microcell-mediated chromosome transfer that forced specific chromosomes upon the cells^10,11^ and might experience accumulation of massive chromosomal rearrangements^12^. To date, no study has systematically profiled the genomic and transcriptomic landscapes of an isogenic aneuploid model, coupling these landscapes with genetic and pharmacological vulnerabilities. Therefore, a system of non-transformed, p53-WT isogenic cells that evolved aneuploidy through chromosome mis-segregation followed by natural selection, could be of high value for studying the molecular and cellular consequences of aneuploidy *per se*.

A major stressful consequence of aneuploidy is genomic instability. Aneuploidy has been associated with increased levels of DNA damage^13^: chromosome segregation errors promote genomic instability via several mechanisms, and aneuploidy itself can lead to perturbed DNA replication, DNA repair and mitosis^14–21^. This association is bi-directional, as replication stress can trigger structural and numerical chromosomal instability (CIN), resulting in aneuploidy^22^. Interestingly, aneuploid cancer cells have been shown to be resistant to DNA damage-inducing agents^7,23–26^, and this increased resistance has been linked to their overall drug resistance^7,24^, to their delayed cell cycle^26^, or to specific protective karyotype alterations^23,25^. Whether the ongoing genomic instability of aneuploid cells leads to elevated DNA damage repair (DDR) activity that could protect them from further induction of DNA damage, has remained an open question. In addition, it is currently unknown whether specific signaling pathways are activated in aneuploid cells in response to the elevated DNA damage, and whether any such pathway might present a therapeutic opportunity.

Here, we established a library of stable RPE1 clones with various degrees of aneuploidy. We performed unbiased genomic and functional characterizations of 6 such isogenic clones and revealed increased vulnerability of aneuploid cells to RAF/MEK/ERK pathway inhibition, and specifically to CRAF perturbation, which could also sensitize cells to DNA damage-inducing chemotherapies. This novel aneuploidy-induced functional dependency was validated in human cancer cell lines, and may therefore be important for the development of novel cancer therapeutics, as well as for improved application of existing anticancer drugs.

## Results

### A novel tool model system to dissect the consequences of aneuploidy on cell physiology

To investigate which pathways are critical for the survival of aneuploid cells, we generated a novel system of isogenic aneuploid cell lines (and matching pseudo-diploid counterparts) derived from the untransformed, pseudo-diploid, immortalized retinal pigment epithelial cell line RPE1-hTERT (henceforth RPE1). This library was generated by transiently treating RPE1 with reversine, a small-molecule inhibitor of the mitotic kinase MPS1, followed by single-cell sorting and clonal expansion^17,27,28^ (**Fig. 1a**) (for more details see **Methods**). Out of an initial pool of ∼5,000 single-cell sorted cells, ∼200 clones (4%) were able to proliferate and were subjected to shallow whole-genome sequencing to determine their karyotypes. This revealed that 79 clones (corresponding to ∼40% of the 200 clones, **Fig. 1b, Supp. Table 1**) showed one or more aneuploid chromosome(s), on top of the gains of chromosome 10q and chromosome 12, which are known to pre-exist in the parental RPE1 cells^7,23^.

**Figure 1.**
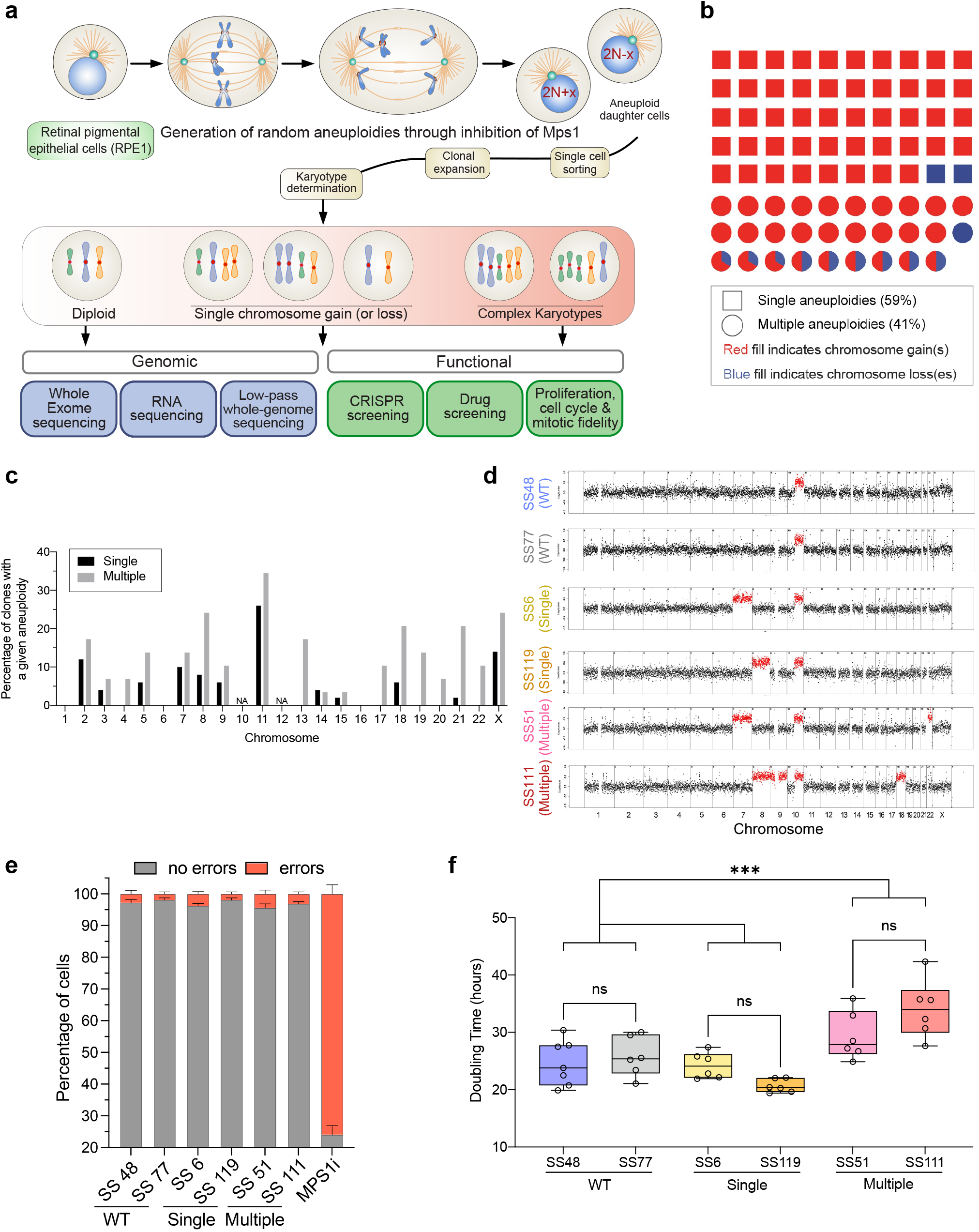
Characterization of matched aneuploid and pseudo-diploid clones. **(a)** Schematic representation of clone generation. See the main text for a detailed description. **(b)** Chart showing the percentage of clones harboring single and multiple aneuploidies. Each shape represents a clone, with 79 clones shown in total. Squares represent clones with single aneuploidies, and circles represent clones with multiple aneuploidies. Red and blue indicate chromosome gains and losses, respectively, and the proportion of each color within the circle represents the fraction of gains/losses out of the aneuploid chromosomes. Clones harboring aneuploidies for chromosomes 10q and 12 were excluded, as they are already abundant in the parental RPE1 population. The library is enriched in clones harboring trisomies over monosomies (****, p<0.0001, Chi-square Goodness of Fit test), and monosomies are more tolerated in multiple aneuploidies background than in single aneuploidy background (**, p=0.0024, Chi-square test). **(c)** Quantification of the percentage of clones harboring a given aneuploid chromosome in single (black) and multiple (grey) aneuploid clones. Chromosomes 10 and 12 were excluded, as a high fraction of the parental RPE1 cells already harbor a gain of one or both of these chromosomes. NA: not applicable. **(d)** Low-pass whole-genome sequencing (lp-WGS) copy number profiles, showing the karyotypes of selected pseudo-diploid (SS48 and SS77) and aneuploid (SS6, SS119, SS51, SS111) clones derived from RPE1 cells. Chromosome gains are colored in red, including the clonal gain of the q-arm of chromosome 10. Resulting karyotype is indicated in brackets. **(e)** Quantification of chromosome segregation errors (including lagging chromosomes, micronuclei, and anaphase bridges) determined by live-cell imaging. Treatment with the MPS1 inhibitor, reversine, was used as positive control. n=4 independent experiments. Graph shows the average ± SEM. **(f)** Doubling time of pseudo-diploid (SS48 and SS77) and aneuploid (SS6, SS119, SS51, SS111) clones. n=7 (SS48) and n=6 (SS77, SS6, SS119, SS51, SS111) independent experiments. n.s., p>0.25; ***, p=0.0005; One-way ANOVA, Tukey’s multiple comparison test.

Among the aneuploid clones present in our library, about 60% displayed single aneuploidies, whereas ∼40% harbored multiple aneuploidies (**Fig. 1b, Extended Data Fig. 1a** and **Supp. Table 1**). Among the clones with single aneuploidies, the vast majority (48 out of 50, 96%) harbored trisomies, and monosomies were present only in a few clones (2 out 50, 4%). Interestingly, chromosome losses were much more common in the context of a complex karyotype: ∼28% (8 out of 29) of clones carrying multiple aneuploidies displayed at least one chromosome loss, a significant enrichment over the monosomy representation in the clones with a single aneuploidy (**Fig. 1b**). Analysis of the identity of aneuploid chromosomes revealed that the frequency at which a given chromosome was gained or lost in stable single-aneuploid clones was below 15% for most chromosomes, with the exception of chromosome 11 that was present in ∼25% of the aneuploid clones (**Fig. 1c**). Further, ∼40% of chromosomes were completely absent from the library of single aneuploidies (chromosomes 1, 4, 6, 13, 16, 17, 19, 20 and 22; **Fig. 1c**), but most of them were gained in clones harboring multiple aneuploidies (with the exception of chromosomes 1, 6 and 16; **Fig. 1c**). Importantly, whole-chromosome aneuploidies were much more common (∼90% of clones) than segmental aneuploidies (∼10%; **Extended Data Fig. 1b**), in line with the known effects of chromosome mis-segregation induced by MPS1 inhibition^17,27,28^, and with previous reports that structural aneuploidies could be tolerated only in a *TP53*-deficient background^29^.

Although our library is not large enough to enable statistical analyses of specific chromosome occurrence and co-occurrence patterns, our data suggest that the absence of some chromosomes from the library is mostly a consequence of selection, rather than of skewed chromosome mis-segregation. Single-cell whole-genome sequencing (scWGS) of parental RPE1 cells^30^ immediately following reversine exposure did not reveal a strong bias in the aneuploidy prevalence across chromosomes (**Extended Data Fig. 1c**), although the mild aneuploidy biases that were observed were highly similar to those recently reported^31^ (higher-than-average aneuploidy rates for chromosomes 1-5, 8, 11 and X; lower-than-average rates for chromosomes 14, 15 and 19-22; **Extended Data Fig. 1c**). However, these mild biases could not explain the chromosome composition bias observed in our final library, which was much more dramatic (with 9 chromosomes not appearing at all as single trisomies). Moreover, the relative aneuploidy prevalence of each chromosome immediately after treatment was not significantly correlated with its library representation. Overall, our analysis shows that randomly generated aneuploidies tend to be detrimental, with single monosomies being less tolerated than trisomies, and with some karyotypes being less fit than others, likely due to selection towards fitter clones.

### Proliferation, mitotic fidelity and cell cycle analyses of the RPE1 clones

For further studies, we decided to focus on aneuploid cell lines either trisomic for a given chromosome or harboring a complex karyotype in which the same chromosome gain was present in combination with other karyotypic alterations. Thus, we selected six clones for further characterization: two pseudo-diploid control clones, RPE1-SS48 and RPE1-SS77 (henceforth SS48 and SS77); two clones with single chromosome gains, RPE1-SS6 and RPE1-SS119 (henceforth SS6 and SS119), which are trisomic for chromosome 7 and 8, respectively; and two clones with complex karyotypes, RPE1-SS51 that is trisomic for chromosomes 7 and 22, and RPE1-SS111 that is trisomic for chromosomes 8, 9 and 18 (henceforth SS51 and SS111) (**Fig. 1d**; note that gain of the q-arm of chromosome 10 is a clonal event in RPE1 cells). In addition, the selected chromosomes cover a large portion of the size and coding spectrum of human chromosomes [Chr7, 159 mega–base pairs (Mbp) and 1,048 coding genes; Chr8, 146 Mbp and 659 coding genes; Chr7 + Chr22, 159 + 50 Mbp and 1,048 + 488 coding genes; Chr8 + Chr9 + Chr18, 146 + 138 + 80 Mbp and 659 + 786 + 270 coding genes].

Having isolated the aneuploid clones, we asked whether their karyotypes were stable, since aneuploidy can often lead to chromosomal instability^16,17,19,21,23,25^. For this, we evaluated the fidelity of chromosome segregation by live cell imaging. We analyzed the presence of mitotic errors, such as lagging chromosomes, anaphase bridges and micronuclei formation. Both WT and aneuploid clones displayed the same basal level of segregation defects (∼2-5%, **Fig. 1e**) and did not show significant differences in mitotic timing, which was ∼25 min for all clones (**Extended Data Fig. 1d**). As positive control for mitotic defects, we treated RPE1 cells with reversine, which led to both chromosome segregation errors and shortening of mitotic timing (**Fig. 1e** and **Extended Data Fig. 1d**; in agreement with^17,27,28^). To further confirm that aneuploid clones remained chromosomally-stable over time, we profiled their karyotypes by low-pass WGS (lp-WGS) following 10 passages in culture, and found that their chromosomal composition has not evolved (**Extended Data Fig. 1e;** compare karyotypes to those shown in **Fig. 1d**). The stability of the aneuploid karyotypes is important as it should allow us to assess the cellular consequences of aneuploidy *per se*, rather than those of the chromosomal instability that is often closely associated with aneuploidy.

Aneuploidy has a detrimental effect on cell cycle progression^7,11,17,28,32–34^. To assess the effect of aneuploidy on cell proliferation, we compared population doublings between pseudo-diploid and aneuploid clones. Our analysis demonstrated that the proliferation rate of clones harboring single trisomies was similar to that of pseudo-diploid clones, displaying a doubling time of roughly 24 hours (**Fig. 1f**), while clones with complex karyotypes displayed a longer population doubling time (SS51 displaying a doubling time of 29 hours and SS111 of 34 hours; **Fig. 1f** and **Extended Data Fig. 1f**). In sum, our efforts led to the generation of a library of matched non-transformed cells with various degrees of aneuploid, stable karyotypes. This library is a powerful tool for the genomic and functional characterization of the consequences of aneuploidy.

### Unbiased genomic and functional characterization of the RPE1 clones

To characterize the genomics of our library of matched non-transformed cells with various degrees of aneuploidy, we performed whole-exome sequencing (WES), and followed standard pipelines for point mutation and copy number calling (**Methods**). In line with RPE1 being a non-transformed cell line, only a handful of cancer-relevant mutations were observed in the clones, the majority of which shared by all clones (**Extended Data Fig. 2a** and **Supp. Table 2**). Surprisingly, however, the analysis revealed that SS77, one of the near-diploid control clones, acquired a clonal heterozygous p53-inactivating mutation (**Extended Data Fig. 2a-b**). WES-based copy number analysis confirmed the karyotypes of the clones (**Supp. Table 3**), in full agreement with the lp-WGS results (**Fig. 1d**). Further analysis revealed that the highly-aneuploid clones, SS51 and SS111, carried many more focal copy number alterations (CNAs), compared to the pseudo-diploid clone SS48 and to the single-trisomy clones, SS6 and SS119 (**Fig. 2a**), suggesting a higher degree of genomic instability in these clones. Consistent with the acquisition of a p53-inactivating mutation, the number of CNAs in the SS77 clone was comparable to that in the highly-aneuploid clones (**Extended Data Fig. 2c**). These results suggested that SS77 should not be used as a pseudo-diploid clone, but could be used instead to confirm that identified vulnerabilities are indeed associated with aneuploidy *per se* and not with the status of p53 pathway activation. Therefore, for downstream validation and mechanistic experiments (as detailed below), we used another *TP53*-WT pseudo-diploid cell line, RPE1-SS31 (henceforth SS31), after confirming that its karyotype and proliferation rate were comparable to those of SS48 (**Extended Data Fig. 3a-e**).

**Figure 2:**
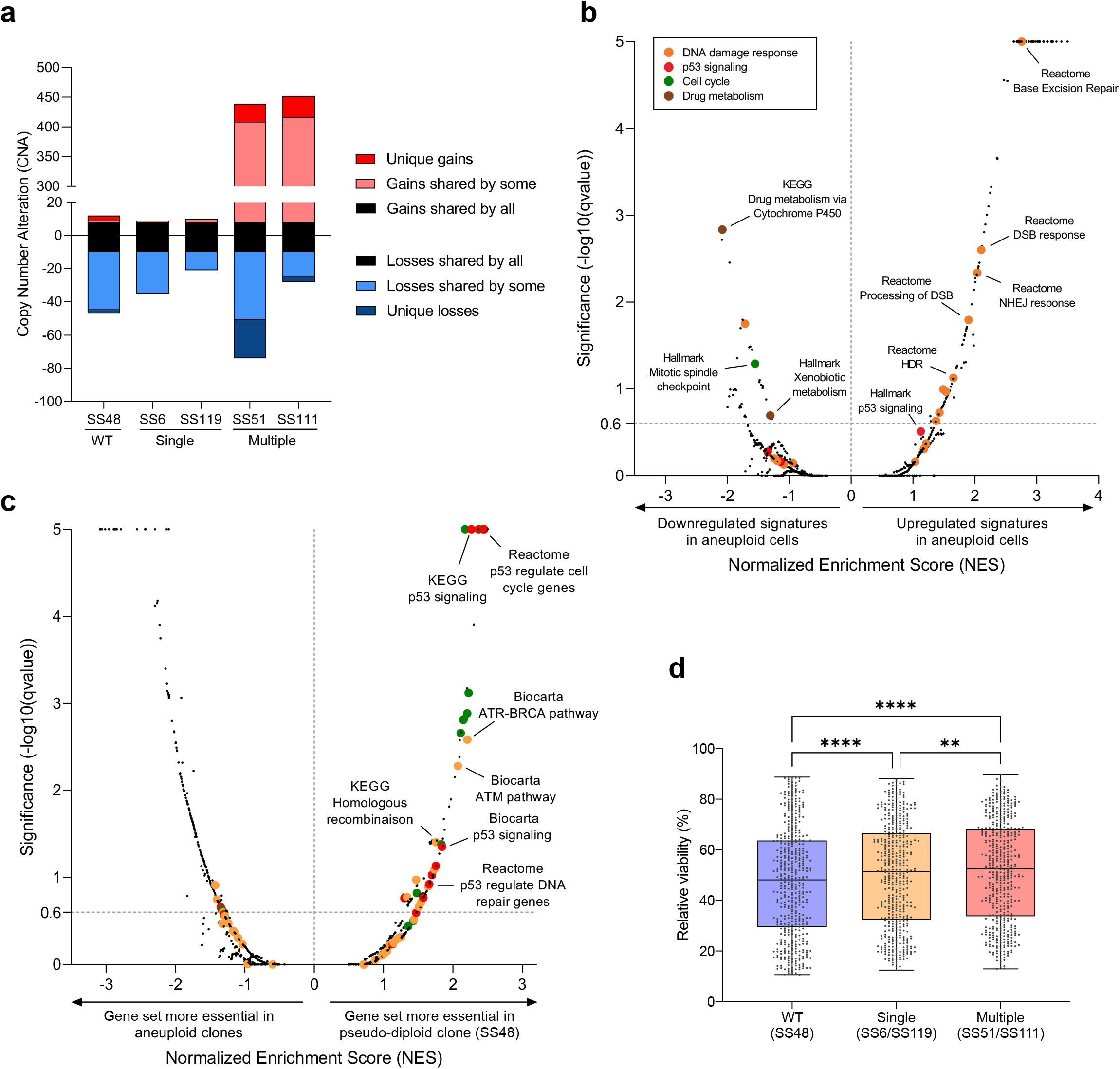
Unbiased genomic and functional characterization of RPE1 clones. **(a)** Copy number alterations (CNAs) across the RPE1 clones. Highly-aneuploid clones, SS51 and SS111, exhibited the highest number of CNAs. **(b)** Comparison of the differential gene expression patterns (pre-ranked GSEA results) between the near-diploid SS48 clone (control) and the highly-aneuploid SS51 and SS111 clones. Plot presents enrichments for the Hallmark, KEGG, Biocarta and Reactome gene sets. Significance threshold set at qvalue=0.25. Enriched pathways are color-coded. **(c)** Comparison of the differential gene dependency scores (pre-ranked GSEA results) between the near-diploid SS48 clone (control) and the aneuploid SS6, SS119 and SS51 clones. Plot presents enrichments for the Hallmark, KEGG, Biocarta and Reactome gene sets. Significance threshold set at qvalue=0.25. Enriched pathways are color-coded. **(d)** Comparison of overall drug sensitivity between a near-diploid control clone (SS48), clones with a single trisomy (SS6 and SS119), and clones with multiple trisomies (RPE1-SS51 and RPE1-SS111). Only drugs that led to a viability reduction ranging from -10% to -90% compared to DMSO control (see **Methods**) were considered (n=456 drugs). **p=0.004, ****p<0.0001; Repeated-Measures One-way ANOVA, Tukey’s multiple comparison test.

We continued the genomic characterization of the clones by investigating their gene expression profiles using RNA sequencing (RNAseq). Principal component analysis (PCA) showed that the highly-aneuploid clones, SS51 and SS111, clustered together despite harboring a completely different set of trisomies (**Extended Data Fig. 2d**). Next, we performed a differential gene expression analysis, followed by pre-ranked gene set enrichment analysis (GSEA^35,36^), to identify gene expression signatures that are induced by aneuploidy regardless of the specific affected chromosome(s) (**Supp. Tables 4-5**). As expected, the over-expressed genes in each aneuploid clone were enriched for the gained chromosome(s) (**Extended Data Fig. 2e**). Importantly, however, chromosome-independent transcriptional signatures could also be identified. The aneuploid clones significantly upregulated signatures related to DNA damage response and repair (DDR) (**Fig. 2b** and **Extended Data Fig. 2f**), suggesting that the aneuploid cells cope with elevated levels of DNA damage, consistent with the CNA analysis (**Fig. 2a**). The aneuploid clones also significantly upregulated signatures related to RNA metabolism and pathways associated with management of proteotoxic stress (**Supp. Tables 4-5**), suggesting altered gene expression process in the aneuploid clones, as detailed in our companion manuscript (Ippolito, Zerbib et al, *bioRxiv* 2023). On the other hand, aneuploid clones significantly down-regulated transcriptional signatures associated with cell cycle (**Fig. 2b**), in line with their slower proliferation rates (**Fig. 1d**), as well as signatures associated with drug metabolism (**Fig. 2b**).

Next, we performed a functional characterization of the sensitivity of the isogenic cell lines to genetic and pharmacological perturbations, in order to obtain an unbiased view of the relationship between aneuploidy and cellular vulnerability. We first performed genome-wide CRISPR/Cas9 screens in the 6 clones (**Methods**). One of the clones, SS111, failed quality control and was therefore excluded from downstream analyses. We calculated the gene dependency scores for 18,120 genes in all RPE1 clones. We then compared the genetic dependencies between the aneuploid clones and the pseudo-diploid SS48 clone, using pre-ranked GSEA, to identify pathways that are preferentially essential either in pseudo-diploid or in aneuploid clones (**Supp. Table 6**). Interestingly, the aneuploid clones were less sensitive than the pseudo-diploid clone to knockout of genes related to DNA damage response (**Fig. 2c**), suggesting that their adaptation to elevated levels of DNA damage may enable them to cope better with further DNA damage induction. Of note, the aneuploid clones were also less sensitive than the pseudo-diploid clone to knockout of genes associated with cell cycle progression/regulation (**Fig. 2c**), in line with their slower proliferation rates (**Fig. 1f**). In addition, blocking p53 activity promoted cell proliferation to a greater extent in the aneuploid clones, reflected in our analysis by a decreased ‘sensitivity’ of the aneuploid clones to p53 pathway perturbation (**Fig. 2c**). These results suggest that the aneuploid clones may already activate the p53 pathway under standard culture conditions. We note that we also found aneuploid cells to be more dependent on mechanisms related to RNA and protein metabolism (**Supp. Table 6**), and followed up on these findings in a companion study (Ippolito, Zerbib et al, *bioRxiv* 2023).

Finally, we performed a pharmacological screen of 5,336 small molecules, using the Broad Drug Repurposing Library^37^, which is composed of preclinical compounds and FDA-approved drugs with known mechanisms of action, allowing further comparison of the vulnerability landscapes of pseudo-diploid vs. aneuploid cells. Each RPE1 clone was exposed to 2.5µM of each compound in duplicates, and cell viability was assessed after 72hrs (**Methods** and **Supp. Table 7**). Interestingly, aneuploid clones were significantly more resistant to drug treatment in general, compared to the pseudo-diploid clone SS48 (**Fig. 2d**), consistent with the observed downregulation of drug metabolism (**Fig. 2b**). The more aneuploid the cells, the more resistant they were to drug treatments, in line with previous reports that linked increased aneuploidy with reduced drug sensitivity^7,24,26^. Importantly, aneuploid cells were also found to be more sensitive to specific classes of drugs (as detailed in the next sections). Notably, the differential vulnerabilities identified in the genetic and pharmacological screens were recapitulated when the p53-mutated, yet chromosomally unaltered, pseudo-diploid clone SS77 was included in the analysis (**Extended Data Fig. 4a-b**, further demonstrating that the identified differential sensitivities are indeed related to the aneuploid state of the cells.

To enable broad use of this new resource by the community, the datasets from the omics profiling and the genetic perturbation screens have been incorporated into the Dependency Map portal (www.depmap.org), and the drug screen results have been incorporated into the Drug Repurposing Hub portal (www.broadinstitute.org/drug-repurposing-hub).

As aneuploid cancer cells have been previously shown to experience DNA damage^13,14^, and to be more resistant to cell cycle inhibitors^26^, to DNA damage inducers^7,23,25^, and to drugs in general^7^, we conclude that our isogenic non-transformed cell line models are capable of capturing aneuploidy-induced cellular consequences that also apply to cancer. Given that increased dependencies of aneuploid cells to these molecular pathways have not been reported before, we decided to focus our downstream validation and mechanistic studies on DDR (the current study) and RNA and protein metabolism (Ippolito, Zerbib et al, *bioRxiv* 2023).

### Elevated DDR and increased resistance of aneuploid cells to DNA damage induction

Highly-aneuploid clones exhibited elevated transcriptional signatures of multiple DNA damage and repair gene sets in comparison to the pseudo-diploid clone SS48 (**Fig. 3a** and **Extended Data Fig. 5a**). To assess DNA damage in the pseudo-diploid and highly-aneuploid clones, we used immunofluorescence microscopy to quantify γH2AX foci in the cells. The number of positive nuclei was significantly higher in the highly-aneuploid clones (**Fig. 3b-c**), consistent with the increased CNA prevalence in these cells (**Fig. 2a**).

**Figure 3:**
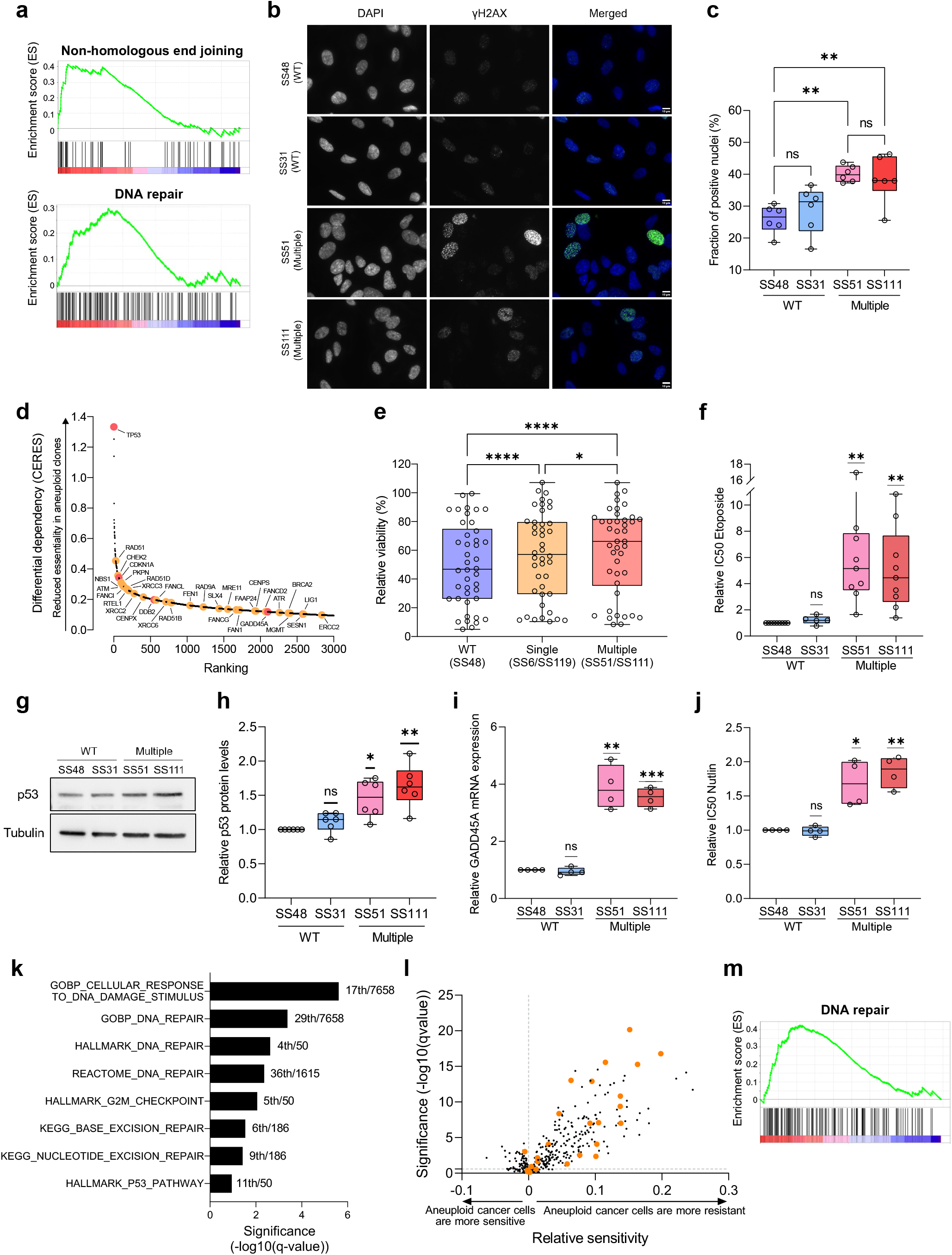
Elevated DDR and increased resistance of aneuploid cells to DNA damage induction. **(a)** Gene set enrichment analysis (GSEA) of DNA damage response (DDR) gene expression signatures, comparing the highly-aneuploid clones, SS51 and SS111, to the pseudo-diploid clone SS48. Shown are enrichment plots for the Reactome ‘Non-Homologous End Joining’ gene set (NES=2.06, q-value=0.0026), and the Hallmark ‘DNA repair’ gene set (NES=1.72, q-value=0.013). **(b)** Immunofluorescence of yH2AX foci in pseudo-diploid clones, SS31 and SS48, and in highly-aneuploid clones, SS51 and SS111. Green, yH2AX; Blue, DAPI; Scale bar, 10μm. **(c)** Quantitative comparison of yH2AX foci between pseudo-diploid clones (SS48 and SS31) and highly-aneuploid clones (SS51 and SS111). n=6 independent experiments; **, p=0.0022 (SS51/SS48) and p=0.0066 (SS111/SS48), *, p=0.0167 (SS51/SS31) and p=0.0462 (SS111/SS31); One-Way ANOVA, Tukey’s multiple comparison. **(d)** The top 3,000 genes that aneuploid clones were most preferentially resistant to their knockout in comparison to the pseudo-diploid clone SS48, based on our genome-wide CRISPR/Cas9 screen. Highlighted in are key genes that belong to the p53 pathway (in red) or to DNA damage response (in orange). **(e)** Comparison of cellular sensitivity to drugs that directly induce DNA damage (alkylating and intercalating agents, anti-topoisomerases and PARP inhibitors) between pseudo-diploid (SS48), single-trisomy clones (SS6 and SS119) and clones with multiple trisomies (SS51 and SS111). Only drugs that led to a viability reduction ranging from -10% to -90% in at least one group were considered (n=42 drugs). *, p=0.0482; ****, p<0.0001; Repeated-Measure One-Way ANOVA, Tukey’s multiple comparison test. **(f)** Comparison of drug sensitivity (determined by IC50 values) to 72hr drug treatment with etoposide, between pseudo-diploid clones (SS48 and SS31) and highly-aneuploid clones (SS51 and SS111). n=5 (SS31) and n=9 (SS48, SS51, SS111) independent experiments. IC50 fold-change was calculated relative to SS48, per experiment. **, p=0.0081 and p=0.0046, for SS51 and SS111, respectively; One-Sample t-test. **(g)** Western blot of p53 protein levels in pseudo-diploid clones (SS48 and SS31) and highly-aneuploid clones (SS51 and SS111). **(h)** Quantification of p53 protein levels, calculated relative to SS48 per experiment. n=6 independent experiments. *, p=0.0104 and **, p=0.0042, for SS51 and SS111 respectively; One-Sample t-test. **(i)** Comparison of the mRNA levels of the p53 transcriptional target GADD45A, quantified by qRT-PCR, between pseudo-diploid clones (SS48 and SS31) and highly-aneuploid clones (SS51 and SS111). n=4 independent experiments. Expression fold-change was calculated relative to SS48, per experiment. **, p=0.0049 and ***, p=0.0006 for SS51 and SS111 respectively; One-Sample t-test. **(j)** Comparison of drug sensitivity (determined by IC50 values) to 72hr drug treatment with nutlin-3a, between pseudo-diploid clones (SS48 and SS31) and highly-aneuploid clones (SS51 and SS111). n=4 independent experiments. IC50 fold-change was calculated relative to SS48, per experiment. *, p=0.0265 and **, p=0.0052 for SS51 and SS111 respectively; One-Sample t-test. **(k)** Gene set enrichment analysis of the genes whose expression correlates with proliferation in highly-aneuploid cancer cell lines but not in near-diploid cancer cell lines. Significant enrichment of multiple DNA repair signatures was observed. The MSigDB ‘Hallmark’, ‘KEGG’, ‘Reactome’, and ‘Gene Ontology-Biological Process’ gene sets were analyzed (separately). Significance values represent the FDR q-values. The ranking of each DDR signature (out of all signatures included in the gene set collection) is indicated next to each bar. **(l)** Differential drug sensitivities between near-euploid and highly-aneuploid human cancer cell lines, based on the large-scale GDSC drug screen^45^. Data are taken from Cohen-Sharir *et al*^7^. Direct DNA damage inducers (alkylating and intercalating agents, anti-topoisomerases and PARP inhibitors) are highlighted in orange. Highly-aneuploid cell lines are more resistant to this class of drugs. (**m**) Pre-ranked GSEA of mRNA expression levels showing that high aneuploidy levels are associated with upregulation of the DNA damage response (DDR) in human primary tumors. Shown is the enrichment plot of Hallmark ‘DNA repair’ (NES=1.73; q-value=0.001) gene set. Data were obtained from the TCGA mRNA expression dataset^86^.

As we found that the aneuploid clones were less sensitive than the pseudo-diploid clone to knockout of genes related to DNA damage response (**Fig. 2d**), we next focused on these genes. This list included important genes involved in the response to both single-strand breaks (SSBs) and double-strand breaks (DSBs), such as *RAD51, CHEK2, ATM, ATR, BRCA2*, and the *MRE11*-*NBN* complex^38,39^ (**Fig. 3d**). This result suggests that the aneuploid clones are more resistant than the pseudo-diploid clones to further induction of DNA damage. We therefore investigated the response of aneuploid clones to small molecules that directly induce DNA damage (a total of 42 compounds in our pharmacological screen). Indeed, aneuploid cells were significantly less sensitive to these drugs, and drug resistance was correlated with the degree of aneuploidy: clones with a single trisomy (SS6 and SS119) were more resistant than the pseudo-diploid clone (SS48), and clones with multiple trisomies (SS51 and SS111) were more resistant than clones with single trisomies (**Fig. 3e**). To validate these results, we treated two pseudo-diploid and two highly-aneuploid clones with two clinically relevant chemotherapies: the topoisomerase II inhibitor etoposide, which induces DSBs, and the topoisomerase I inhibitor topotecan, which induces SSBs. The highly-aneuploid clones were significantly more resistant to both drugs compared to pseudo-diploid clones (**Fig. 3f** and **Extended Data Fig. 5b**).

*TP53* came up as the most differentially essential gene in the pseudo-diploid clones (**Fig. 3d**). A plausible explanation for this would be that the p53 pathway is already active in the aneuploid clones, so that the proliferation boost provided by its inhibition is greater in these cells. Indeed, GSEA analysis of the gene expression profiles revealed significant up-regulation of p53 targets in the aneuploid clones (**Extended Data Fig. 5c**), consistent with the increased DNA damage observed in these clones. Western blot confirmed elevated levels of the p53 protein in the highly-aneuploid clones relative to their pseudo-diploid counterparts (**Fig. 3g-h**). Moreover, qRT-PCR analysis of transcriptional downstream targets of p53 identified increased expression of several p53 targets, including those specifically linked to the DNA damage response, such as GADD45A, a key player in the G2/M DNA damage checkpoint^40^ (**Fig. 3i** and **Extended Data Fig. 5d**). Finally, we treated the RPE1 clones with nutlin-3a, an inhibitor of the MDM2-p53 interaction causing p53 stabilization, and found that the highly-aneuploid clones were significantly more resistant to p53 activation than the pseudo-diploid clones (**Fig. 3j**). Together, these findings suggest that the aneuploid clones experience higher DNA damage levels, leading to p53 pathway activation and increased DDR, which render them less sensitive to further induction of DNA damage (or to further p53 activation).

To assess the generalizability of these findings, we turned to a second isogenic system of RPE1 cells and their aneuploid derivatives, RPTs^8^. In this system, inhibition of cytokinesis led to tetraploidization of the RPE1 cells, resulting in chromosomal instability that soon made them highly-aneuploid^8^. γH2AX staining revealed significantly more ongoing DNA damage in the aneuploid RPT cells in comparison to their pseudo-diploid parental cells (**Extended Data Fig. 5e-f**). Moreover, the RPT cells were more resistant to both etoposide and topotecan (**Extended Data Fig. 5g-h**), and their resistance patterns matched their pre-existing DNA damage levels. Interestingly, RPT3 is the most aneuploid clone of this system^8^, and we found it to exhibit the highest degree of DNA damage and to be the most resistant to further DNA damage induction. Therefore, the increased DNA damage and the subsequent resistance to DNA damage induction characterized the aneuploid cells in two independent RPE1-based cellular systems in which aneuploidy was induced in completely different ways.

Lastly, we addressed whether these findings also apply to aneuploid human cancer cells. We extended our published table of aneuploidy scores of human cancer cell lines^7^ to ∼1,500 cancer cell lines (**Supp. Table 8** and **Methods**). Matched doubling times were available for ∼500 of these cancer cell lines^41^, allowing us to investigate whether DDR is required for the proliferation of aneuploid cells. Specifically, we found that the genes associated with the proliferation capacity of highly-aneuploid, but not of near-euploid, cell lines were enriched for DDR signatures (**Methods, Supp. Table 9, Fig. 3k**). Moreover, aneuploid human cancer cells were significantly more resistant to chemical agents that directly induce DNA damage across several independent drug screens^42–45^ (**Fig. 3l** and **Extended Data Fig. 5i-j**). Finally, a lineage-controlled pancancer analysis of The Cancer Genome Atlas (TCGA) showed a significant elevation of the DDR gene expression signature in highly-aneuploid human tumors (**Fig. 3m** and **Extended Data Fig. 5k**). We conclude that ongoing DNA damage, activated DDR, and increased resistance to DNA damage induction, are fundamental characteristics of both non-transformed and cancerous aneuploid cells.

### Increased CRAF activity and dependency in aneuploid cells

We next analyzed our pharmacological screen to identify increased vulnerabilities of the aneuploid clones. Although the aneuploid clones were generally more resistant than the pseudo-diploid clones to drug treatments (**Fig. 2e**), they were significantly more sensitive to RAF/MEK/ERK pathway inhibition (**Fig. 4a**). We validated the differential drug sensitivity to two of the top differentially-active drugs, TAK632 and 8-Br-cAMP, and found that the highly-aneuploid clones were significantly more sensitive to both of these RAF inhibitors (**Fig. 4b-c**). Interestingly, TAK632 is a pan-RAF inhibitor exhibiting increased affinity for CRAF (also known as Raf-1) over BRAF^46^, and 8-Br-cAMP was previously described as a specific CRAF inhibitor^47^, suggesting a specific role of CRAF in the observed RAF dependency. Previous studies have shown that CRAF activation follows BRAF/CRAF heterodimerization^48,49^. Thus, we treated the cells with PLX7904, a novel RAF inhibitor developed to inhibit BRAF/CRAF heterodimerization and the resultant CRAF activation^50,51^. Consistent with the response to the other two RAF inhibitors, highly-aneuploid clones were more sensitive to PLX7904 compared to pseudo-diploid clones (**Extended Data Fig. 6a**), validating the dependency of highly-aneuploid clones on CRAF activity.

**Figure 4:**
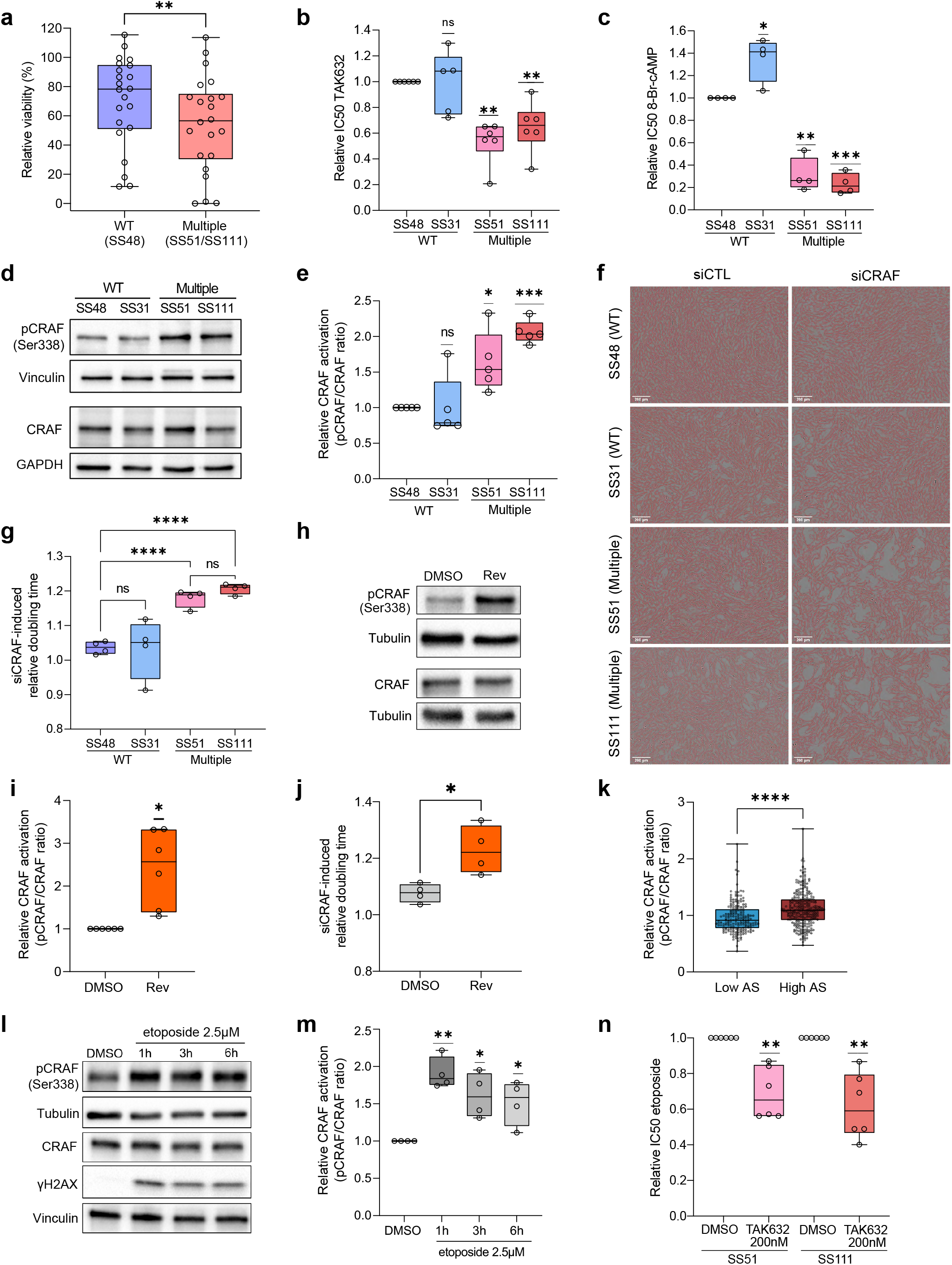
Aneuploid cells exhibit increased activity and dependency to CRAF, which is functionally linked to DNA damage repair. **(a)** Comparison of the sensitivity to RAF/MEK/ERK inhibitors (n=22 drugs) between the near-diploid control clone (SS48) and the highly-aneuploid clones (SS51 and SS111). Only drugs that led to a viability reduction ranging from -10% to -90% in at least one of the groups were considered. **, p=0.0018, two-tailed paired t-test. **(b-c)** Comparison of drug sensitivity (determined by IC50 values) to 72hr drug treatment with the CRAF inhibitors TAK632 (**b**) and 8-Br-cAMP (**c**), between pseudo-diploid clones (SS48 and SS31) and highly-aneuploid clones (SS51 and SS111). IC50 fold-change was calculated relative to SS48, per experiment. TAK632: n=5 (SS31) and n=6 (SS48, SS51, SS111) independent experiments; **, p=0.001 and p=0.007, for SS51 and SS111, respectively. 8-Br-cAMP: n=4 independent experiments; **, p=0.0017, ***, p=0.002, for SS51 and SS111, respectively; One-Sample t-test. **(d)** Western blot of pCRAF (Ser338) and total CRAF protein levels in pseudo-diploid clones (SS48 and SS31) and highly-aneuploid clones (SS51 and SS111). Vinculin and GAPDH were used as housekeeping control. **(e)** Quantification of CRAF activation, based on the pCRAF/CRAF ratio in each clone, calculated relative to SS48 per experiment. n=5 independent experiments. *, p=0.028; ***, p=0.001, for SS51 and SS111 respectively; One-Sample t-test. **(f)** Representative images of pseudo-diploid clones (SS48 and SS31) and highly-aneuploid clones (SS51 and SS111) treated with an siRNA against CRAF. The cytostatic effect of the knockdown was stronger in the highly-aneuploid clones. Cell masking (shown in red) was performed using Ilastik for visualization purposes. Scale bar, 200µM. **(g)** Doubling time quantification in the pseudo-diploid clones (SS48 and SS31) and highly-aneuploid clones (SS51 and SS111) treated with an siRNA against CRAF. Proliferation rate was calculated relative to a control siRNA treatment, per experiment. n=4 independent experiments. ****, p<0.0001 (SS51/SS48, SS111/SS48, SS51/SS31, SS111/SS31); One Way ANOVA, Tukey’s multiple comparison. **(h)** Western blot of pCRAF and total CRAF protein levels in parental RPE1 cells treated with reversine (500nM) or control DMSO for 20hrs to induce aneuploidy, then harvested 72hrs post wash-out. Tubulin was used as housekeeping control. **(i)** Quantification of CRAF activation based on the pCRAF/CRAF ratio in the reversine-treated cells, calculated relative to the DMSO control per experiment. n=6 independent experiments. * p=0.0122; One Sample t-test. **(j)** Comparison of doubling time following siRNA against CRAF, between aneuploidy-induced parental RPE1 cells (treated for 20hr with the SAC inhibitor reversine (500nM)) or with control DMSO, then harvested 72hrs post wash-out. Proliferation rate was calculated relative to a control siRNA treatment, per experiment. n=4 independent experiments; *, p=0.0157; two-tailed unpaired t-test. **(k)** Comparison of CRAF activity, as measured by the pCRAF/CRAF ratio, between the top and bottom aneuploidy quartiles of human cancer cell lines (n=455 cell lines). Data were obtained from the DepMap portal 22Q1 release^57^. CRAF activity level is significantly higher in the highly-aneuploid cancer cell lines. ****, p<0.0001, two-tailed Mann-Whitney test. **(l)** Western blot of pCRAF (Ser338), total CRAF, and γH2AX protein levels in parental RPE1 cells treated with etoposide (2.5µM) for 1, 3 or 6 hours. CRAF activation levels are associated with the degree of DNA damage in the cells. Tubulin and Vinculin were used as housekeeping controls. **(m)** Quantification of CRAF activation based on the pCRAF/CRAF ratio in the etoposide-treated cells, calculated relative to the DMSO control per experiment. CRAF activation levels were highest at 1hr post-etoposide treatment, and declined over time, following γH2AX expression levels. n=4 independent experiments. **, p=0.0034 *, p=0.0267 and p=0.0411, for 1h, 3h and 6h etoposide treatment, respectively; One Sample t-test. **(n)** Comparison of drug sensitivity (determined by IC50 values) to 72hr treatment with the DNA damage inducer etoposide, between highly-aneuploid RPE1 clones (SS51 and SS111) treated with a sub-lethal dose (200nM) of the CRAF inhibitor TAK632, or with DMSO control. CRAF inhibition sensitized the cells to etoposide. n=6 independent experiments. IC50 fold-change was calculated relative to the DMSO-treated cells, per experiment. **, p=0.0033 and p=0.0039, for SS51 and SS111, respectively; One-Sample t-test.

Several studies have pointed to a connection between RAF activity and aneuploidy induction^52–55^. Thus, we measured RAF activation in our clones, using qRT-PCR and WB. BRAF expression levels were consistently elevated only in SS51 (**Extended Data Fig. 6b-d**), which harbors an extra copy of chromosome 7 on which BRAF resides (BRAF is constitutively phosphorylated^56^ so its activity cannot be assessed by measuring phosphorylation). In contrast, CRAF was consistently activated (as measured by pCRAF/CRAF protein ratio) in both highly-aneuploid clones (**Fig. 4d-e**). Together, these results suggest that CRAF is indeed activated in the aneuploid clones, underlying the increased sensitivity of the aneuploid cells to RAF inhibitors. To validate the specific dependency of aneuploid cells on CRAF, we knocked it down using siRNAs (**Extended Data Fig. 6e**). Indeed, CRAF knockdown had an inhibitory effect on the proliferation of highly-aneuploid clones, but not on pseudo-diploid clones (**Fig. 4f-g**). To further validate and characterize the effect of CRAF inhibition, we treated the cells with 8-Br-cAMP and followed their morphology, motility and proliferation using live-cell imaging (**Extended Data Fig. 6f**). The effects of CRAF inhibition on cell proliferation (**Extended Data Fig. 6g-h**), cell morphology ((**Extended Data Fig. 6i-j**), and cell motility (**Extended Data Fig. 6k-l**) were all significantly stronger in the highly-aneuploid clones. Finally, we quantified cell death following CRAF inhibition in the clones using flow cytometry, and found that there was no significant difference in cell death between the pseudo-diploid and highly-aneuploid clones (**Extended Data Fig. 6m-n**). We therefore conclude that highly-aneuploid clones preferentially depend on CRAF activity for their proliferation, and that CRAF inhibition is mostly cytostatic, rather than cytotoxic, for the aneuploid cells.

To assess whether CRAF activation is an immediate adaptation of cells following aneuploidy induction, we quantified its activity immediately after reversine treatment. We found increased CRAF activity following MPS1 inhibition both in the parental RPE1 cells (**Fig. 4h-i**) and in the pseudo-diploid clones SS48 and SS31 (**Extended Data Fig. 7a-b**). CRAF activation following reversine-mediated aneuploidization was p53-independent, as we also observed significant increase in CRAF activation upon MPS1 inhibition in *TP53*-KD and *TP53*-KO RPE1 cells (**Extended Data Fig. 7c-f** and **Extended Data Fig. 8**). Importantly, the inhibitory effect of CRAF knockdown on cell proliferation significantly increased following reversine-induced aneuploidization (**Fig. 4j**), confirming that aneuploidy increases the cellular sensitivity to CRAF inhibition. Elevated CRAF activity and increased vulnerability to CRAF inhibition were also recapitulated in the second isogenic system of RPE1 cells and their highly-aneuploid RPT derivatives: the aneuploid RPT clones exhibited elevated levels of both pCRAF and total CRAF levels (albeit not of pCRAF/CRAF protein ratio) (**Extended Data Fig. 7g-j**), and were significantly more sensitive to CRAF inhibition (**Extended Data Fig. 7k**).

We then asked whether CRAF activity is associated with a high degree of aneuploidy in human cancer cells as well. Quantification of the pCRAF/CRAF protein ratio across 455 highly-aneuploid vs. near-euploid cancer cell lines^57^, revealed increased CRAF activity in highly-aneuploid cancer cells (**Fig. 4k**; BRAF and CRAF total protein levels were not changed^58^ (**Extended Data Fig. 7l-m**)). We conclude that aneuploid cells activate CRAF in the context of cancer cells as well.

Interestingly, several studies have recently shown that CRAF activity is functionally linked to DDR^59,60^. Specifically, CRAF was shown to be activated in response to DNA damage, and its pharmacological or genetic inhibition sensitized cells to ionizing radiation or genotoxic drugs^59^. In line with these findings, etoposide treatment in the parental RPE1 cells led to a significant increase in their CRAF activity (**Fig. 4l-m**). To investigate if the increased activation of CRAF in the aneuploid cells is causally related to their increased resistance to DNA damage induction, we treated the highly-aneuploid clones SS51 and SS111 with a sub-lethal dose of the CRAF inhibitor TAK632 for 72hr (**Extended Data Fig. 7n**), in combination with the DSB-inducing drug etoposide. CRAF inhibition sensitized the aneuploid cells to etoposide (**Fig. 4n**), confirming that CRAF activation was required to overcome DNA damage in the aneuploid clones.

### Increased MEK/ERK activity and dependency in aneuploid cells

Having found increased CRAF activity and dependency in aneuploid cells, we next investigated the activation of the canonical CRAF downstream targets, MEK and ERK. Indeed, both MEK and ERK activity was significantly higher in the highly-aneuploid clones than in the pseudo-diploid ones (**Fig. 5a-d**). Together with the increased CRAF activity, these results indicate that the entire RAF/MEK/ERK signaling cascade is elevated in the aneuploid cells. We therefore compared the vulnerability of pseudo-diploid and aneuploid cells to MEK and ERK inhibition. The highly-aneuploid clones were significantly more sensitive to the clinically-approved MEK inhibitors, trametinib and selumetinib (**Fig. 5e** and **Extended Data Fig. 9a**), and to the ERK inhibitor, ulixertinib (also known as BVD-523) (**Fig. 5f**). Together with their increased dependency on CRAF, these results show that aneuploid clones depend on the entire RAF/MEK/ERK pathway.

**Figure 5:**
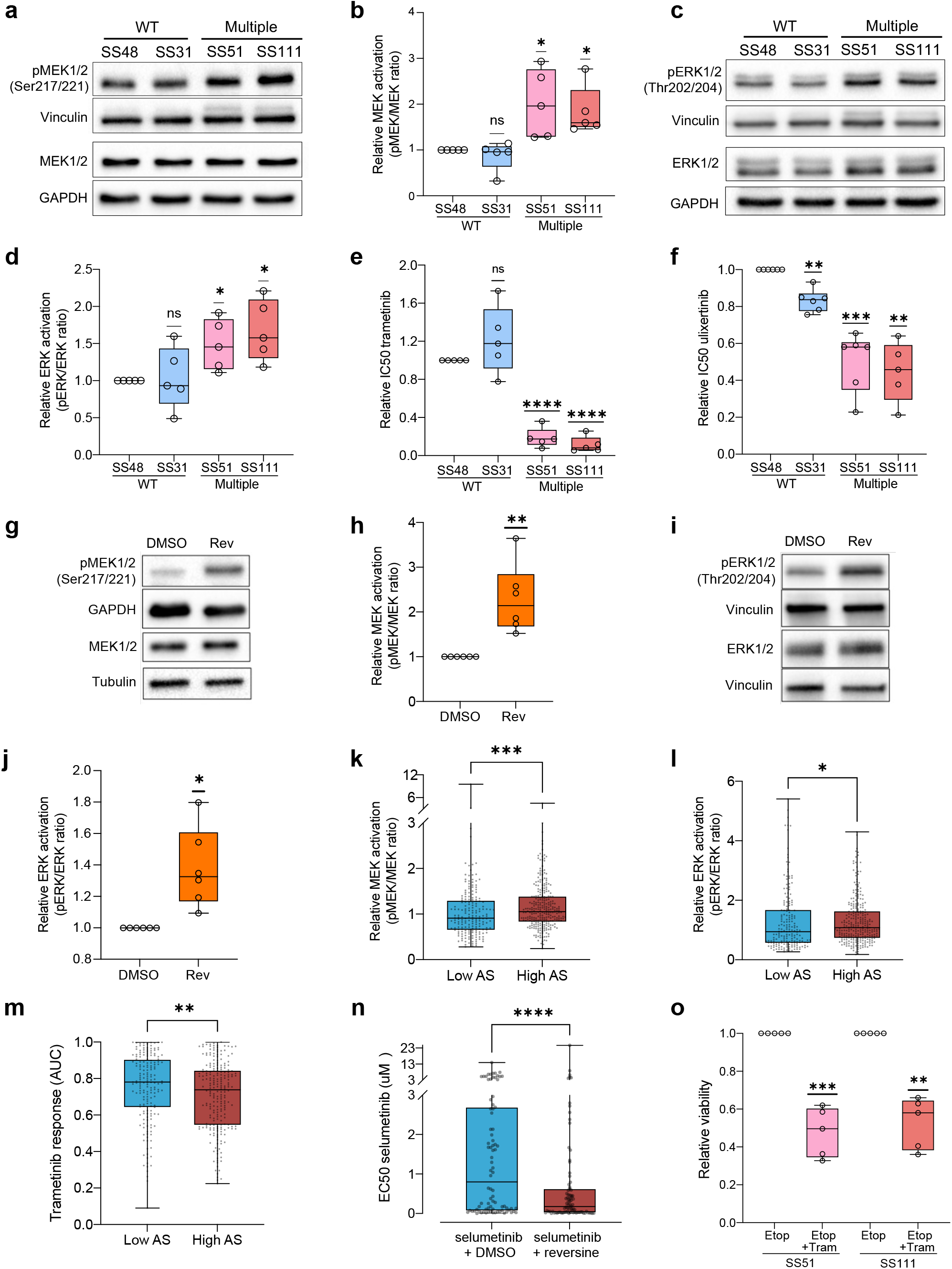
Increased MEK/ERK pathway activity and dependency in aneuploid cells. **(a)** Western Blot of pMEK1/2 (Ser217/221) and MEK1/2 protein levels in pseudo-diploid clones (SS48 and SS31) and highly-aneuploid clones (SS51 and SS111). Vinculin and GAPDH were used as housekeeping controls. **(b)** Quantification of MEK1/2 activation based on the pMEK/MEK ratio in each clone, calculated relative to SS48 per experiment. n=5 independent experiments; *, p=0.0383, p=0.0247, for SS51 and SS111, respectively; One Sample t-test. **(c)** Western Blot of pERK1/2 (Thr202/Tyr204) and ERK1/2 protein levels in pseudo-diploid clones (SS48 and SS31) and highly-aneuploid clones (SS51 and SS111). Vinculin and GAPDH were used as housekeeping controls. **(d)** Quantification of ERK1/2 activation based on the pERK/ERK ratio in each clone, calculated relative to SS48 per experiment. n=5 independent experiments; *, p=0.0346 and p=0.0223 for SS51 and SS111, respectively; One Sample t-test. **(e)** Comparison of drug sensitivity (determined by IC50 values) to 72hr drug treatment with the MEK inhibitor trametinib, between pseudo-diploid clones (SS48 and SS31) and highly-aneuploid clones (SS51 and SS111). IC50 fold-change was calculated relative to SS48, per experiment. n=5 independent experiments; ****, p<0.0001; One-Sample t-test. **(f)** Comparison of drug sensitivity (determined by IC50 values) to 72hr drug treatment with the ERK inhibitor ulixertinib, between pseudo-diploid clones (SS48 and SS31) and highly-aneuploid clones (SS51 and SS111). IC50 fold-change was calculated relative to SS48, per experiment. n=5 independent experiments; **, p=0.0011 and p=0.0016, ***, p=0.0007 for SS31, SS111 and SS51, respectively; One-Sample t-test. **(g)** Western blot of pMEK1/2 (Ser217/221) and total MEK1/2 protein levels in parental RPE1 cells treated with reversine (500nM) or with control DMSO for 20hrs to induce aneuploidy, then harvested 72hrs post wash-out. Tubulin and GAPDH were used as housekeeping control. **(h)** Quantification of MEK activation based on the pMEK/MEK ratio in the reversine-treated cells, calculated relative to the DMSO control per experiment. n=6 independent experiments. **, p=0.0096; One-Sample t-test. **(i)** Western blot of pERK1/2 (Thr202/Tyr204) and total ERK1/2 protein levels in parental RPE1 cells treated with reversine (500nM) or with control DMSO for 20hrs to induce aneuploidy, then harvested 72hrs post wash-out. Vinculin was used as housekeeping control. **(j)** Quantification of ERK activation based on the pERK/ERK ratio in the reversine-treated cells, calculated relative to the DMSO control per experiment. n=6 independent experiments. *, p=0.0148; One-Sample t-test. **(k)** Comparison of MEK activity, as measured by the pMEK/MEK ratio, between the top and bottom aneuploidy quartiles of human cancer cell lines (n=460 cell lines). Data were obtained from the DepMap portal 22Q1 release^57^. MEK activity levels are significantly higher in the highly-aneuploid cancer cell lines. ***, p=0.0006; two-tailed Mann-Whitney test. **(l)** Comparison of ERK activity, as measured by the pERK/ERK ratio, between the top and bottom aneuploidy quartiles of human cancer cell lines (n=460 cell lines). Data were obtained from the DepMap portal 22Q1 release^57^. ERK activity levels are significantly higher in the highly-aneuploid cancer cell lines. *, p=0.0424; two-tailed Mann-Whitney test. **(m)** Comparison of drug sensitivity (determined by AUC) to the MEK inhibitor trametinib, between the top and bottom aneuploidy quartiles of human cancer cell lines (n=412 cell lines). Data were obtained from GDSC1 drug screen, DepMap portal 22Q1 release. **, p=0.0069; two-tailed Mann-Whitney test. **(n)** PRISM-based comparison of drug sensitivity (determined by EC50 values) to 120hr treatment with the MEK inhibitor selumetinib, between cancer cells treated with the SAC inhibitor reversine (250nM) or with control DMSO (n= 84 cell lines). Aneuploidy induction sensitized cancer cells to selumetinib. ****, p<0.0001; two-tailed Wilcoxon rank sum test. **(o)** Comparison of viability following 72hrs treatment with a sub-lethal dose (0.45nM) of the MEK inhibitor trametinib or DMSO, in combination of DNA damage induction using Etoposide (2.5uM) in highly-aneuploid RPE1 clones (SS51 and SS111). MEK inhibition sensitized highly aneuploid clones to DNA damage induction. n=5 independent experiments. Fold change in viability after combination was calculated relative to etoposide-treated cells, per experiment; ***, p=0.0009 and **, p=0.0015, for SS51 and SS111 respectively; One-Sample t-test.

To assess whether MEK and ERK activation is an immediate response of cells to aneuploidy induction, we quantified their activity after treating the cells with reversine. MEK and ERK activities both increased significantly following MPS1 inhibition (**Fig. 5g-j**). MEK and ERK activation following reversine-mediated aneuploidization was p53-independent, as we also observed significant increase in pathway activation in *TP53*-KD and *TP53*-KO RPE1 cells (**Extended Data Fig. 9b-i**) We then examined whether MEK and ERK activities are also associated with a high degree of aneuploidy in human cancer cells. Quantification of pMEK/MEK and pERK/ERK protein ratio across hundreds of highly-aneuploid vs. near-euploid cancer cell lines^57^ revealed the increased activity of both MEK and ERK in highly-aneuploid cancer cells (**Fig. 5k-l**), consistent with the increased CRAF activity **(Fig. 4k**). Thus, we conclude that the increased activity of the RAF/MEK/ERK pathway is associated with a high degree of aneuploidy in cancer cells as well.

The sensitivity of aneuploid cells to MEK inhibitors is of particular importance given their clinical use. Comparison of the drug response of near-euploid and highly-aneuploid human cancer cell lines to trametinib and selumetinib revealed that aneuploid cancer cells were significantly more sensitive to these MEK inhibitors (**Fig. 5m** and **Extended Data Fig. 9j**). To confirm that MEK dependency is indeed causally related to aneuploidy in cancer cells, we performed a pooled screen of cell lines, using the PRISM barcoded cell line platform^43^, and assessed the response of 578 human cancer cell lines to selumetinib, in combination with a low dose (250nM) of reversine or with a vehicle-control (**Methods**). Whereas the proliferation effect of reversine itself at this low concentration was mild (**Supp. Table 10**), it significantly sensitized the cancer cell lines to MEK inhibition (**Fig. 5n**). We thus conclude that highly-aneuploid cancer cells are more sensitive to MEK inhibition.

Several studies have documented a beneficial effect of combining MEK/ERK inhibitors with DDR inhibitors in multiple myeloma and pancreatic cancer^61,62^. As we found that CRAF inhibition could sensitize aneuploid cells to DNA damage induction^59,60^ (**Fig. 4l-n**), we next investigated whether MEK inhibition could also sensitize aneuploid cells to DNA damage induction. A combination of a sub-lethal dose of trametinib (**Extended Data Fig. 9k**) significantly sensitized highly-aneuploid clones to etoposide (**Fig. 5o** and **Extended Data Fig. 9l**), consistent with the role of the CRAF/MEK/ERK pathway in overcoming DNA damage.

Therefore, we propose that aneuploid cells increase their CRAF/MEK/ERK pathway activity, which helps them overcome the increased amount of DNA damage. Inhibition of MEK/ERK signaling could therefore sensitize aneuploid cells to DNA damage inducers.

## Discussion

Aneuploidy has been recognized as a pervasive feature of tumors for over 100 years^63^. Early last century, it was proposed that the presence of unbalanced chromosome numbers is a hallmark of cancer cells. More recently, efforts employing sequencing technologies have confirmed that virtually all tumors harbor karyotypic abnormalities, spanning from segmental to whole-chromosomal aneuploidies^6^. Nevertheless, research on aneuploidy has been hampered by the paucity of suitable systems to model it *in vitro* and, more importantly, by the inability to disentangle aneuploidy from other features often co-existing in cancer cells, such as p53 inactivation and genomic instability. Thus, studying and understanding the effects of karyotypic abnormalities on cell physiology while controlling for potential confounders remains of paramount importance for cancer biology. Likewise, deconstructing the pathways deregulated by the aneuploid state – and understanding the cellular mechanisms employed by cancer cells to cope with aneuploidy-induced cellular stresses – holds the promise of unraveling novel and unique dependencies exploitable for cancer therapy^7^.

To investigate the cellular and molecular consequences of aneuploidy, we have generated, characterized and analyzed a library of untransformed human cell lines with stable and defined aneuploid karyotypes. We employed multiple genomic, transcriptomic and functional assays to extensively profile this new isogenic cell line library (**Fig. 1**,**2**), and have incorporated these data sets into the Dependency Map (www.depmap.org) and the Drug Repurposing Hub (www.broadinstitute.org/drug-repurposing-hub), in order to enable broad use of this new resource by the community. Our own functional analyses and validation experiments revealed that aneuploid cells have increased activation of DDR and RNA metabolism, resulting in altered dependencies of aneuploid cells on these pathways.

### Increased dependency on RAF/MEK/ERK pathway activity in aneuploid cells

Aneuploidy has been previously reported to directly correlate with increased levels of DNA double-strand breaks, mutational loads^19,20^ and replication stress^15–17,20–22,29,64^. Here, by using our system of matched aneuploid cells, complemented by additional isogenic systems of aneuploidy and comprehensive analyses of human cancer cell lines, we found that aneuploid cells are more resistant to DNA damage inducers and to DDR perturbation in general (**Fig. 3**), in line with previous reports of increased resistance of aneuploid cancer cells to DNA damage-inducing drugs^7,24,26,65,66^. Our findings uncover the pathways triggered in response to DNA damage, and highlight the importance of RAF/MEK/ERK pathway activity, and CRAF in particular, in this regard (**Fig. 4**). CRAF has been implicated in DNA damage response through both kinase-dependent and kinase-independent mechanisms. CRAF kinase activity can directly feed into the RAF/MEK/ERK pathway to ensure proper execution of DDR^48,49,67^. Inhibitors of the RAF/MEK/ERK pathway have been recently reported to increase cellular dependency on functional DDR^61,62,68–70^. In agreement with this notion, our data points at a DDR regulation of RAF/MEK/ERK pathway in aneuploid cells, enabling aneuploid cells to tolerate DNA damage and to keep proliferating in its presence (**Fig. 5**). These findings raise the exciting possibility to combine clinically-approved RAF/MEK/ERK inhibitors with DNA damage-inducing chemotherapies for the targeting of aneuploid tumors.

Notably, the finding that aneuploidy activates the RAF/MEK/ERK pathway has broader implications that go beyond the interaction of this pathway with the DDR. Indeed, activation of RAF/MEK/ERK pathway is observed in ∼40% of tumors^71^ and is the consequence of oncogenic mutations of different molecular players operating in this signaling cascade. Mutations in RAS and RAF genes – and of CRAF in particular – have been associated with high degree of CIN and aneuploidy^54,72–75^, lending further support to the importance of this pathway for the cellular response to aneuploidy. Notably, RPE1 cells are KRAS-mutant, but our findings clearly indicate that the pathway does not reach its maximum activity in the parental population and is further activated following aneuploidy induction. Therefore, the finding that the aneuploid state *per se* leads to increased CRAF and RAF/MEK/ERK pathway activities – and to increased vulnerability to perturbation of this pathway – independently of co-occurring mutations, indicates that aneuploid tumors may benefit from treatment with RAF/MEK/ERK inhibitors regardless of genetic mutations in this pathway.

Several kinase-independent roles of CRAF have been reported as well, mainly relying on its scaffolding functions^59,76–80^. For example, CRAF has been shown to be pivotal in supporting the activation of CHK2, a crucial player in DDR^59^. Thus, it remains formally possible that the CRAF dependency observed in aneuploid clones might also be the consequence of CHK2 activation mediated by kinase-independent roles of CRAF. Intriguingly, however, our analysis found that aneuploid clones were less sensitive to *CHEK2* knockout yet more dependent on CRAF activity, suggesting that CHK2 activity might be largely dispensable in aneuploid cells. CRAF also plays a role in regulating Aurora B, PLK1 and Aurora A^81,82^, crucial components of the mitotic spindle, which act to ensure proper chromosome segregation. Therefore, CRAF perturbation may result in DNA damage accumulated during aberrant mitoses. Future studies will be aimed at fully dissecting CRAF mode of action in response to DNA damage in aneuploid cells and exploring if other unknown mechanisms operate to sense and respond to DNA damage following aneuploidization.

### RAF/MEK/ERK pathway activity and p53 activation

The p53 pathway is a major barrier for aneuploidy tolerance^2,6,14,29^. We and others have shown that aneuploidy-associated stresses can actively lead to p53 activation: oxidative, metabolic, genotoxic and proteotoxic stresses can lead to increased p53 pathway activity followed by cell cycle arrest and reduced proliferation capabilities of aneuploid cells^11,16,17,21,28,29,83^. Among these stresses, aneuploidy-associated DNA damage can instigate p53 activation in several ways, including: lagging chromosomes that get broken by the cleavage furrow during the process of mis-segregation^14^, ruptured micronuclei exposing their DNA to cytoplasmic nucleases^84,85^, segmental chromosomes generated as a result of aneuploidy-induced genome instability and DNA replication stress^16,17,21,29^. In agreement with these data, our aneuploid clones show increased signs of DNA damage, high levels of p53 expression and upregulation of its target genes compared to pseudo-diploid counterparts (**Fig. 3**).

Notably, although p53 activation and aneuploidy-induced stresses are intimately intertwined, we found increased dependency on the RAF/MEK/ERK pathway (**Fig. 4, 5**) independently of p53 status. Indeed, although we discovered these dependencies in p53-WT cells, these effects remained significant when: (a) the aneuploid cells were compared to a near-diploid control clone harboring a p53-inactivating mutation (SS77) (**Extended Data Fig. 2, 4**); (b) aneuploidy was induced on the background of p53 knock-down or knock-out (**Extended Data Fig. 7-9**); and (c) hundreds of human cancer cell lines – most of them p53-inactivated – were stratified based on their aneuploidy scores, showing a positive correlation between the degree of aneuploidy and the activation and dependency on RAF/MEK/ERK (**Fig. 4, 5**). We conclude that the vulnerabilities identified through our functional analyses are indeed a consequence of the aneuploid state *per se*. We note that our functional studies focused on aneuploid cells with extra chromosomes (trisomies), which is characteristic of most human tumors^6,7^. As the consequences of monosomy may differ from those of trisomy^9^, future studies should examine the relevance of these functional dependencies to tumors that are predominantly affected by chromosome losses.

### Concluding remarks

Extensive DNA damage is one of the most prominent consequences of aneuploidy. Our work points at the central role of the RAF/MEK/ERK pathway in overcoming aneuploidy-induced DNA damage, enabling cells to tolerate this major aneuploidy-induced stress (**Fig. 6**). Our findings may have important implications for the selective targeting of aneuploid cancer cells by perturbing these pathways: selective inhibition of the RAF/MEK/ERK pathway, and of CRAF in particular, might sensitize aneuploid cancer cells to treatments with DNA damage inducers. If these unique cellular vulnerabilities of aneuploidy hold true in the clinical setting, we speculate that they could be exploited for the selective eradication of aneuploid tumors.

**Figure 6:**
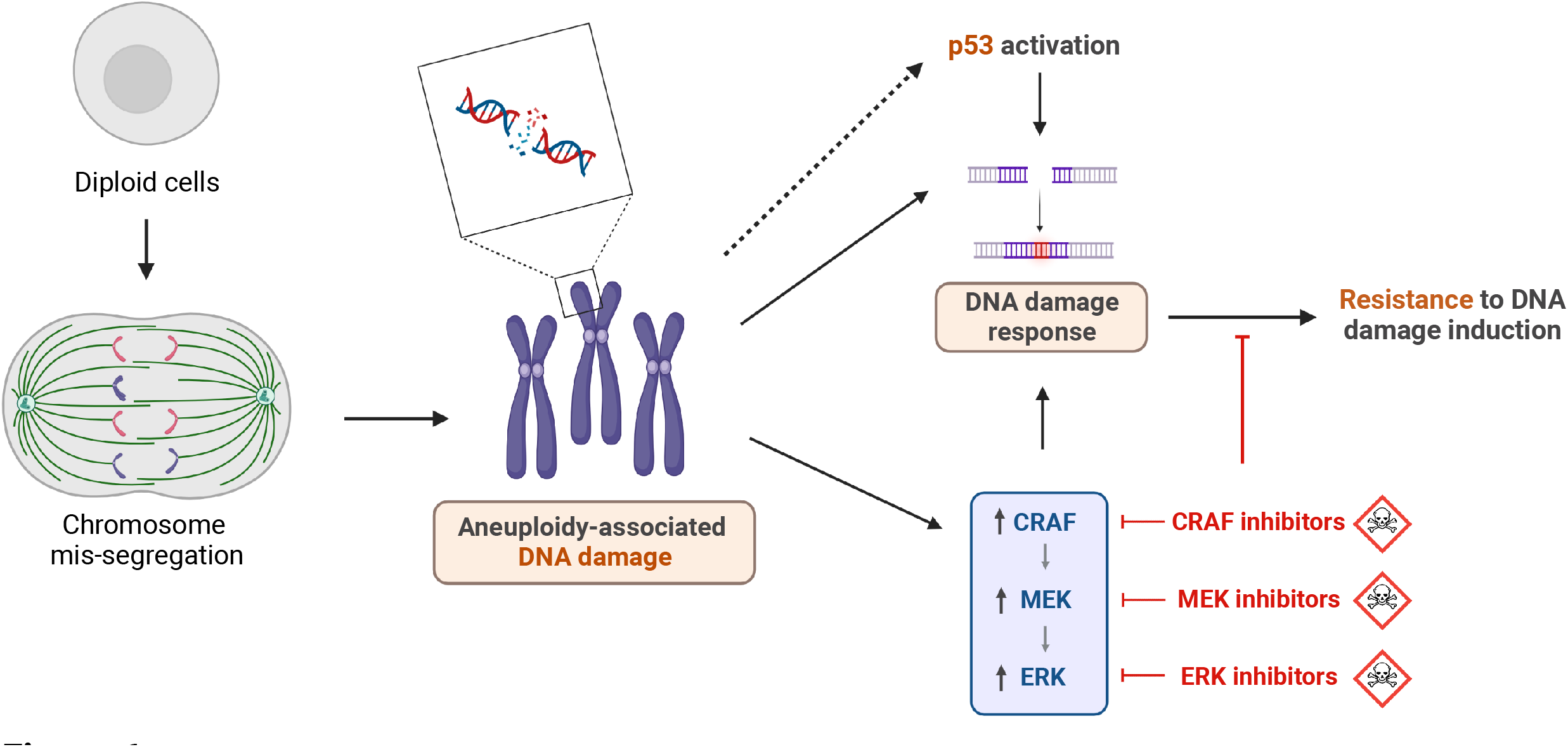
Aneuploidy-induced DNA damage results in the upregulation of the CRAF/MEK/ERK pathway. A summary illustration of the study. When cells become aneuploid following chromosome mis-segregation, they acquire DNA damage that activates the DNA damage response (DDR). When p53 is intact, this results in p53 pathway activation. The increased basal levels of DDR render the cells more resistant to further induction of DNA damage. In parallel, acquisition of DNA damage activates the CRAF/MEK/ERK pathway, which fuels the DNA damage response. Consequently, aneuploid cells are more dependent than their diploid counterparts on CRAF, MEK, and ERK. Pharmacological induction of DNA damage further increases both the DNA damage response and the activation of CRAF/MEK/ERK pathway, and pharmacological inhibition of the CRAF/MEK/ERK pathway can thus sensitize aneuploid cells to DNA damage-inducing chemotherapies.

## Methods

### Cell culture

RPE1-hTERT cells (female cell line, RRID: CVCL_4388), their derivatives clones and RPT were cultured in DMEM (Life Technologies) with 10% fetal bovine serum (Sigma-Aldrich), 1% sodium pyruvate, 4mM glutamine, 100 U/ml penicillin and 100 μg/ml streptomycin. Cells were cultured at 37°C with 5% CO2 and are maintained in culture for maximum three weeks. All cell lines were tested free of mycoplasma contamination using Myco Alert (Lonza, Walkersville, MD, USA) according to the manufacturer’s protocol. To induce random aneuploidy, cells were seeded and synchronized with 5mM Thymidine for 24hrs, then treated with 500nM reversine (or vehicle control) for 16hrs. Read-outs were performed 72hrs post reversine wash-out.

### Generation of a library of aneuploid clones

RPE1-hTERT cells were seeded in 10 cm dishes and treated with 500nM reversine (or vehicle control) for 24 hours. After drug (or vehicle control) wash-out, cells were kept in culture for 2 weeks and split regularly to keep them at about 70/80% confluence. Cells were then trypsinized and single-cell sorted in ∼5000 well of multi-well plates containing conditioned medium (half of the final volume of the well). Single clones were then monitored over a month. Those able to proliferate over this period were transferred into 96 well plates and further expanded to 48, 24, 12 and 6 well plates. Clones were then transferred into 10 cm dishes and further propagated.

### Cell proliferation Assay

RPE1-hTERT derived clones were plated in a 24well plate support in at least three technical replicates. Cells were pictured every 4 hours until reaching confluence using the Incucyte (Satorius). To estimate the confluency, the Built-In program (2021A version) was used applying a threshold of 1 and a minimum area of 140um2 to exclude the debris. Based on these proliferative curves, doubling time was calculated.

### Video microscopy

Live cell imaging was performed using an inverted microscope (Nikon Eclipse Ti) with a 20x objective. The microscope was equipped with an incubation chamber maintained at 37°C with 5% CO2. For experiments shown in **Figure 1** and **Extended Data Figure 1**, RPE1-hTERT derived clones expressing a GFP-tagged version of H2b were seeded on 12-well plates. Cells were filmed for 72h every 5 minutes. For the positive control, cells were immediately treated with DMSO or reversine 500nM. 80 cells for mitotic timing and 60 cells for chromosome segregation fidelity, both from four biological replicates, were analyzed using FIJI software.

### Whole Exome Sequencing and data analysis

WES data were generated as previously described^57^. Mutation calling was performed as previously described^57^, and is available on DepMap (21Q3 release). Heterozygous TP53 mutation was visualized using the Integrative Genomics Viewer (https://software.broadinstitute.org/software/igv/). Copy number calling was performed as previously described^57^ and is available on DepMap 21Q3 release (https://figshare.com/articles/dataset/DepMap_21Q3_Public/15160110). CNAs were defined as copy number values that deviated away of the chromosome-mean CNA value by >0.1 (log2CN) and >5SD (to remove noise, SD calculation excluded deviations >0.24 away of the basal ploidy).

### RNAseq and data analysis

RNA was extracted in triplicates from each of the clones and the quality was assessed using Bioanalyzer 2100. For each sample, RNA library was prepared using TruSeq Stranded total RNA kit (Illumina) following manufacturer’s protocol, and sequenced using TruSeq RNA UDIndices adaptors (Illumina) on Novaseq 6000 sequencer (Illumina) following manufacturer’s protocol. RNA sequence reads were aligned to the human reference genome hg38 using Bowtie2. Normalized read counts, PCA analysis, and differential gene expression analysis were generated using DESeq2 R package^87^. Genes with fewer than 10 normalized read counts were excluded from further analyses. A pre-ranked GSEA was performed on the differentially expressed genes using GSEA software 4.0.3, with the following parameters: 1000 permutations and Collapse analysis, using the Hallmark, KEGG, Biocarta, and Reactome gene sets (in separate analyses). For the pre-ranked GSEA analysis, genes with fewer than 20 normalized read counts were excluded.

### Genome-wide CRISPR screens and data analysis

Cells were barcoded and treated as previously described^88^. CRISPR dependency scores (CERES scores) were calculated as previously described^88^ and are available on DepMap 21Q3 release (https://figshare.com/articles/dataset/DepMap_21Q3_Public/15160110). A pre-ranked GSEA was performed on the differentially-expressed genes using GSEA software 4.0.3, with the following parameters: 1000 permutations and Collapse analysis, using the Hallmark, KEGG, Biocarta, and Reactome gene sets (in separate analyses).

### Pharmacological screens and data analysis

Cells were screened against the Drug Repurposing Library from the Broad Institute^37^, as previously described^43^. Briefly, cells were seeded using a Multidrop™ Combi Reagent Dispenser (ThermoFisher) in a 384-well plate, 300 cells per well, in duplicate. 5,336 compounds were tested at 2.5μM. All compounds were pre-plated onto the assay plates prior to cell addition using the Beckman Coulter Labcyte Echo. 72hr post-treatment, cell viability was assessed by CellTiterGlo® (Promega). The viability effect of each compound was calculated for each clone, and compared between the aneuploidy groups (RPE1-SS48 and RPE1-SS77 as near-diploid control clones, RPE1-SS6 and RPE1-SS119 as clones with single trisomies, RPE1-SS51 and RPE1-SS111 as clones with multiple trisomies). The percent activity of each compound was determined by averaging the normalized activity of both replicates. The normalized activity was determined by the following equation –

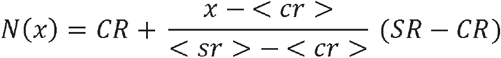

where N is the normalized activity value, x is the measured raw signal of a well, <cr> is the median of the measured signal values of the Central Reference (DMSO control), <sr> is the median of the measured signal values of the Scale Reference (Inhibitor control), CR is the desired median normalized value for the Central Reference (0), and SR is the desired median normalized value for the Scale Reference (−100). Genedata Screener and Spotfire were used in activity normalizations and hit calling. The activity threshold was set at the (negative) of three times the standard deviation of the DMSO control, the direction corresponding to activation or inhibition. Each compound was given one of three designations depending on their activity for each replicate. Compounds were classified as “Active” if the mean of both replicates was equal or less than the activity threshold. Compounds were classified as “Inconclusive” if one of the two replicates was equal or less than the activity threshold but the mean of both replicates was above the activity threshold. Compounds were classified as “Inactive” if neither of the replicates was equal or less than the activity threshold. Only drugs that led to a viability reduction ranging from -10% to -90% in all clones were considered. For comparisons of drugs targeting a specific pathway, a less stringent criterion was applied, so that only drugs that led to a viability reduction ranging from -10% to - 90% in at least one category of cell lines were considered.

### Drug treatments

2,000 cells per well were seeded in a 96w plate using Multidrop™ Combi Reagent Dispenser (ThermoFisher). 24hrs later, cells were treated with drugs of interest. Cell viability was measured after 72hrs (or at the indicated time point) using the MTT assay (Sigma M2128), with 500ug/mL salt diluted in complete medium and incubated at 37°C for 3 hrs. Formazan crystals were extracted using 10% Triton X-100 and 0.1N HCl in isopropanol, and color absorption was quantified at 570nm and 630nm. Absolute IC50 for each drug was calculated using GraphPad PRISM 9.1, inhibitor vs. normalized response (four parameters) equation. All drugs details are available in **Supp. Table 11**.

To test whether CRAF or MEK inhibition sensitized cells to DNA damage induction, 2,000 cells per well of the highly-aneuploid clones RPE1-SS51 and RPE1-SS111 were seeded in triplicates in 96-well plates. Cells were treated with serial dilutions of etoposide in combination with 200nM TAK632 (or vehicle control), or 0.45nM trametinib (or vehicle control) in combination with 2.5uM etoposide, for 72hrs. Cell viability was measured using the MTT assay (Sigma M2128).

### Immunofluorescence

Cells were washed with PBS and then fixed for 15min at room temperature (RT) with 4% para-formaldehyde, followed by permeabilization with Triton X-100 0.5% for 5min at RT, and quenching reduction with L-Glycin 0.1M in PBS for 15min at RT. Slides were then blocked for 30min at RT in blocking solution containing 10% goat serum, 3% BSA, L-Glycin 1%, NaCl 150mM, TRIS pH7.5 10mM, and 0.1% Triton X-100. Slides were incubated with primary antibody against phospho-histone Ser139 γH2AX (1:1000, Millipore) in blocking solution for 1.5hrs at RT in a humid chamber. After washing with PBS, cells were incubated with Alexa Fluor 488 or Alexa Fluor 555 tagged anti-mouse antibody (1:1000, Cell Signaling Technologies) for 1hr at RT in a humid black chamber, and then stained with DAPI (1ug/mL) diluted in PBS for 3min at RT in a humid black chamber. Images were acquired using cellSens Imaging Software (Olympus), and merged using ImageJ. Nuclei containing >5 visible γH2AX foci were considered to be γH2AX-positive. Only cells at interphase were included in the quantification.

### Western blots

Cells were lysed in NP-40 lysis buffer (1% NP-40;150mM NaCl; 50mM Tris HCl pH 8.0) with the addition of protease inhibitor cocktail (Sigma-Aldrich #P8340) and phosphatase inhibitor cocktail (Sigma Aldrich #P0044). Protein lysates were sonicated (Biorector) for 5min (30sec on/30sec off) at 4°c, then centrifuged at maximum speed for 15 min and resolved on 12% SDS-PAGE gels. Bands were detected using chemiluminescence (Millipore #WBLUR0500) on Fusion FX gel-doc (Vilber). All antibodies details are available in **Supp. Table 11**.

### qRT-PCR

Cells were harvested using Bio-TRI® (Bio-Lab) and RNA was extracted following manufacturer’s protocol. cDNA was amplified using GoScript™ Reverse Transcription System (Promega) following manufacturer’s protocol. qRT-PCR was performed using Sybr® green, and quantification was performed using the ΔCT method. All primer sequences are available in **Supp. Table 11**.

### Dependency Map data analysis

Aneuploidy scores (AS) of each cell line were assigned following similar principles to those used by Cohen-Sharir *et al*^7^. Briefly, the median relative copy number was calculated per chromosome arm, the variation across chromosome arms was evaluated, and the number of chromosome arms that deviate from the basal ploidy was determined as the aneuploidy score. The resultant aneuploidy score list is available in **Supp. Table 8**. mRNA gene expression values, protein expression values, CRISPR and RNAi dependency scores (Chronos and DEMETER2 scores, respectively) were obtained from DepMap 22Q1 release (https://figshare.com/articles/dataset/DepMap_22Q1_Public/19139906), and compared between the bottom (AS≤8) and top (AS≥21) aneuploidy quartiles.

For doubling time analyses, the doubling time (DT) of each cell line was assigned as previously published^41^. mRNA expression values were floored to log2(TPM+1)=0.1. Within the bottom quartile (AS≤8) and the top quartile (AS≥21), DT was correlated to gene expression utilizing a linear model (lm function in R studio v4.1.1, with lineage as a covariate, using the equation: gene∼DT+lineage), following the method of Taylor *et al*^6^. Genes were determined as overexpressed in highly proliferative aneuploid cancer cells if they were significantly associated with DT within the top AS quartile but not within the bottom AS quartile. Significance thresholds: (log10(p-value)≥2.5) OR (–log10(p-value)≥1.3 AND correlation coefficient<-0.005). The resultant list of genes is available in Supp. Table 8. This list was subjected to gene set enrichment analysis using the ‘Hallmark’, ‘KEGG’, ‘Reactome’ and ‘Gene Ontology Biological Processes’ gene set collections from MSigDB (http://www.gsea-msigdb.org/gsea/msigdb/)^36,89^. Analysis of CRAF, MEK and ERK protein activity was performed by measuring the ratio between the phosphor-protein to the total protein levels, based on an RPPA protein array^58^. Quantification of total proteins was based on the DepMap proteomics data^58^.

### TCGA data analysis

TCGA data were retrieved using TCGAbiolinks R package^86^. Aneuploidy scores (AS) were obtained from Taylor *et al*^6^, and correlated to tumor gene expression using lineage as a covariate (lm function in R studio v4.1.1, using the equation: gene∼AS+lineage), as previously described^6^. Genes were ranked based on their aneuploidy score coefficient, and then subjected to pre-ranked gene set enrichment analysis^36^ using the ‘Hallmark’, ‘Biocarta’, ‘KEGG’, and ‘Reactome’ gene set collections from MSigDB.

### siRNA transfection

Cells were transfected with siRNAs against CRAF (ONTARGETplus SMART-POOL®, Dharmacon), or with a control siRNA (ONTARGETplus SMART-POOL®, Dharmacon) using Dharmafect1 (Dharmacon) following manufacturers’ protocols. To test whether aneuploidy induction sensitized cells to CRAF, cells were seeded and synchronized with Thymidine 5mM for 24hrs, then treated with reversine 500nM for 20hrs. After the reversine pulse, cells were reverse transfected with siRNA against CRAF using Lipofectamine® RNAiMAX (Invitrogen) following the manufacturer’s protocol. Cell growth following siRNA transfection was followed by live cell imaging using Incucyte® (Satorius). The effect on proliferation was calculated by comparing the fold-change of doubling time of the cells in the targeted siRNA vs. control siRNA wells at 72h post-transfection. For visualization, the cell borders were highlighted using AI-trained Ilastik® software.

### Live cell imaging using LiveCyte®

2,000 cells were seeded in triplicates in microscopy-compatible 96-well plates (Corning), and were treated for 72hr with 10µM of 8-Br-cAMP. Cells were imaged every 20min for 72hr using LiveCyte® (Phase Focus), with an inverted microscope at 10X objective (microscope placed in an incubation chamber maintained at 37°C with 5% CO2). Images were acquired using the LiveCyte acquisition software, and single-cell tracking, segmentation and analyses was performed using the LiveCyte analysis software (Phase Focus). Cell doubling time, dry mass doubling time, cellular area and perimeter, instantaneous velocity and track speed were calculated by the automatic LiveCyte® analysis software (Phase Focus).

### Flow cytometry analysis

100,000 cells were seeded in a 6-well plate and treated for 72hr with 10µM of 8-Br-cAMP, and with Etoposide 2.5µM for 72hrs as a positive control. Cells were stained with SYTOX™ Green Ready Flow™ Reagent (Invitrogen), following the manufacturer’s protocol. Flow cytometry acquisition was performed using CytoFLEX® (Beckman Coulter) and data analysis was performed using Kaluza Analysis software 2.1 (Beckman Coulter). The gating of living cells and singlets was common in all the analyzed samples, per experiment. Gating of positive cells (defined as upper half of the pick in etoposide-treated cells) was defined per cell line.

### PRISM screen

PRISM screen was performed as previously described^7,43^. Briefly, cells were plated in triplicate in 384-well plates at 1,250 cells per well. Cells were treated with the MEK inhibitor selumetinib (8 concentrations of threefold dilutions, ranging from 0.9nM to 20µM) in presence of reversine (250nM) or DMSO for 5 days. Cells were then lysed, and lysate plates were pooled for amplification and barcode measurement. Viability values were calculated by taking the median fluorescence intensity of beads corresponding to each cell line barcode, and normalizing them by the median of DMSO control. Dose-response curves and EC50 values were calculated by fitting four-parameter curves to viability data for each cell line, using the R drc package^90^, fixing the upper asymptote of the logistic curves to 1. EC50 comparisons were performed on the 84 cell lines for which well-fit curves (r^2^>0.3) were generated.

### Statistical analyses

The number of cells used for each experiment is available in the method section. Western Blot quantifications were performed using ImageJ®. The numbers of independent experiments and analyzed cell lines of each computational analysis are available in the figure legends. Statistical analyses were performed using GraphPad PRISM® 9.1. Details of each statistical test are indicated in the figure legends. In each presented box plot, the internal bar represents the median of the distribution. In Figures 1E, S3B, S10B and S10E, the bar represents the mean and SEM. Significance thresholds were defined as p-value = 0.05 and q-value = 0.25.

## Acknowledgments

The authors would like to thank James McFarland for bioinformatic support; Gil Ast, Marina Mapelli, Zuzana Tothova and members of the Ben-David and Santaguida labs for helpful discussions; Varda Wexler for assistance with Figure preparation; Zuzana Storchova for providing the RPE1/RPT cell lines; Nicholas Lyons, Jordan Bryan, Samantha Bender and Jennifer Roth for their assistance with the PRISM screen; Kevin Langley and the PhaseFocus team for their assistance with the LiveCyte® Analysis software. We thank the Broad Institute Genomic Perturbation Platform for their assistance with the CRISPR/Cas9 screens, and the Center for the Development of Therapeutics and Repurposing Hub at the Broad Institute for providing the compound library (https://clue.io/repurposing). This work was supported by the European Research Council Starting Grant (grant #945674 to U.B.-D.), the Israel Cancer Research Fund Gesher Award (U.B.-D.), the Azrieli Foundation Faculty Fellowship (U.B.-D.), the DoD CDMRP Career Development Award (grant #CA191148 to U.B.-D.), the Israel Science Foundation (grant #1339/18 to U.B.-D.), the BSF project grant (grant #2019228 to U.B.-D.), the Italian Association for Cancer Research (AIRC-MFAG 2018 - ID. 21665 project to S.S.), Ricerca Finalizzata (GR-2018-12367077 to S.S.), Fondazione Cariplo (S.S.), the Rita-Levi Montalcini program from MIUR (to S.S.) and the Italian Ministry of Health with Ricerca Corrente and 5×1000 funds (S.S.). U.B.-D. is an EMBO Young Investigator. J.Z. was supported by a fellowship of the Israeli Ministry for Immigrant Absorption and by travel awards from the TAU Constantiner Institute and Cancer Biology Research Center. M.R.I. is supported by an AIRC Fellowship (ID 26738-2021). J.Z, Y.E, and G.L. are PhD and MD-PhD students within the graduate school of the Faculty of Medicine, Tel Aviv University. M.R.I., S.M, S.V. and S.S. are PhD students within the European School of Molecular Medicine (SEMM).

## Extended Data Figures Legends

**Extended Data Figure 1: Characterization of the matched aneuploid and pseudo-diploid clones (related to Fig.1)**

**(a)** Karyotypes of the 79 RPE1 aneuploid clones, derived by transient reversine treatment of the parental RPE1 population. Each row represents one single cell-derived clone. **(b)** Quantification of the percentage of aneuploid clones harboring whole (black) or segmental (gray) aneuploidies. Chromosomes 10 and 12 were excluded, as trisomy of chromosome 12 and gain of chromosome 10q already exist in the parental RPE1 population. There are no differences in segmental and whole chromosome aneuploidy composition between clones with single vs. multiple aneuploidies (p=0.61, Fisher’s exact test). **(c)** Aneuploidy scores (defined as log2(CNV) of each chromosome) of RPE1 cells immediately following aneuploidy induction using the MPS1 inhibitor, reversine. Dashed line indicates the average copy number variation across all chromosomes. Chromosomes 10 and 12 were excluded, as trisomy of chromosome 12 and gain of chromosome 10q already exist in the parental RPE1 population. n=3 independent experiments. NA: not applicable. No significant differences were found across all chromosomes (Kruskal-Wallis test, Dunn’s multiple comparisons), and no correlation was found between chromosomal enrichment following reversine pulse and the final library chromosomal enrichment (rho=0.18, p=0.42; Spearman’s correlation). **(d)** Quantification of mitotic timing of pseudo-diploid (SS48, SS77 and SS31) and aneuploid (SS6, SS119, SS51, SS111) clones. Mitotic timings were determined by live-cell imaging of clones that stably express green fluorescent protein fused to histone H2B (H2B-GFP). Mitotic timing was measured from nuclear envelope breakdown to anaphase onset. Treatment with reversine was used as positive control. n=3 (SS48, SS111) or n=4 (SS77, SS6, SS119, SS51, MPS1i) independent experiments. n.s., p>0.05; ***, p<0.001; One-way ANOVA, Tukey’s multiple comparison test. **(e)** Low-pass whole-genome sequencing (lp-WGS) copy number profiles, showing the karyotypes of pseudo-diploid (SS48 and SS77) and aneuploid (SS6, SS119, SS51, SS111) clones derived from RPE-1 cells after 10 passages in culture. Chromosome gains are colored in red, including the clonal gain of the q-arm of chromosome 10. **(f)** Representative proliferation curves of SS48, SS77, SS6, SS119, SS51 and SS111. Relative confluency was estimated every 4hrs during 48hrs. n=5 independent experiments.

**Extended Data Figure 2: Unbiased genomic characterization of RPE1 clones (related to Fig. 2)**

**(a)** Mutation profiles across the RPE1 clones. Shown are only known COSMIC/TCGA hotspot mutations. Note that SS77 acquired a heterozygous *TP53*-inactivating mutation (p.H193N). **(b)** IGV-based visualization of the reads from the *TP53* locus of SS48 and SS77, demonstrating the acquisition of a clonal (AF∼0.5) heterozygous p53-inactivating mutation (p.H193N) in SS77 clone. **(c)** Copy number alterations (CNAs) across the RPE1 clones, including SS77. Note that SS77 shares an elevated number of CNAs with the highly-aneuploid clones. **(d)** Principal Component Analysis (PCA) of the genome-wide gene expression profiles of the RPE1 clones. PC1 and PC2 explain 60% and 18% of the variance between the samples, respectively. Notably, SS51, SS111 and SS77 cluster together. **(e)** Gene Set Enrichment Analysis (GSEA) plots, demonstrating that the overexpressed genes in each RPE1 clone are enriched to the specific chromosome(s) that are gained in that clone. Plots present enrichments for the chromosome-level positional gene sets, based on the comparison between the control clone (SS48) and the various aneuploid clones (SS6, SS119, SS51, SS111). Enrichments scores : SS6 (chr7 : NES=2.00, qvalue<0.0001), SS119 (chr8 : NES=2.5, qvalue<0.0001), SS51 (chr7 : NES=2.42, qvalue<0.0001 ; chr22 : NES=3.88, qvalue<0.0001), SS111 (chr8 : NES=1.92, qvalue=0.0065 ; chr9 : NES=2.04, qvalue=0.0055 ; chr18Q11 : NES=1.22, qvalue=0.1988). **(f)** Comparison of the differential gene expression patterns (pre-ranked GSEA results) between the near-diploid SS48 clone (control) and the aneuploid SS6, SS119, SS51 and S111 clones. Plot presents enrichments for the Hallmark, KEGG, Biocarta and Reactome gene sets. Significance threshold set at qvalue=0.25. Enriched pathways are color-coded.

**Extended Data Figure 3: Characterization of pseudo-diploid clone SS31 (related to Fig.1 and Fig.2)**

**(a)** Low-pass whole-genome sequencing (lp-WGS) copy number profile, showing the karyotype of the pseudo-diploid SS31 clone, derived from RPE1 cells. Chromosome gains are colored in red, including the clonal gain of the q-arm of chromosome 10. **(b)** Quantification of chromosome segregation errors as in Figure 1E. The graph shows the same value shown in Figure 1E, with the addition of clone SS31. Graph shows the average of four biological replicates ± SEM. **(c)** Quantification of mitotic timing of pseudo-diploid (SS48, SS77 and SS31) and aneuploid (SS6, SS119, SS51, SS111) clones. Mitotic timings were determined by live-cell imaging of clones that stably express green fluorescent protein fused to histone H2B (H2B-GFP). Mitotic timing was measured from nuclear envelope breakdown to anaphase onset. Treatment with reversine, the MPS1 inhibitor, was used as positive control. The graph shows the same value shown in Figure S1D, with the addition of clone SS31. n.s., p>0.05; One-way ANOVA, followed by Tukey’s multiple comparison test. **(d)** Doubling time quantification as in Figure 1F, including SS31. N=7 (SS48) and n=6 (SS77, SS31, SS6, SS119, SS51, SS111) independent experiments. n.s., p>0.25; * p=0.032; **, p=0.003; One-way ANOVA, Tukey’s multiple comparison. **(e)** Representative proliferation curves of SS48, SS77, SS31, SS6, SS119, SS51 and SS111. Relative confluency was estimated every 4hrs during 48hrs. The proliferation curves show the same data as in Figure S1F, with the addition of clone SS31.

**Extended Data Figure 4: Unbiased functional characterization of RPE1 clones (related to Fig. 2)**

**(a)** Comparison of the differential gene dependency scores (pre-ranked GSEA results) between the near-diploid SS48 and SS77 clones (control) and the aneuploid SS6, SS119 and SS51 clones. Plot presents enrichments for the Hallmark, KEGG, Biocarta and Reactome gene sets. Significance threshold set at qvalue=0.25. Enriched pathways are color-coded. **(b)** Comparison of overall drug sensitivity between the near-diploid control clone (SS48 and SS77), clones with a single trisomy (SS6 and SS119), and clones with multiple trisomies (SS51 and SS111). Only drugs that led to a viability reduction ranging from -10% to -90% compared to DMSO control (see **Methods**) were considered (n=439 drugs). *, p=0.0119 (Single/WT), ****p<0.0001 (Multiple/WT), **, p=0.0032 (Single/Multiple); Repeated-Measures One-Way ANOVA, Tukey’s multiple comparison.

**Extended Data Figure 5: Increased DDR in response to aneuploidy (related to Fig. 3)**

**(a)** Gene set enrichment analysis (GSEA) of DNA damage response (DDR) gene expression signatures, comparing the highly-aneuploid clones, SS51 and SS111, to the pseudo-diploid clone SS48. Shown are enrichment plots for the Reactome ‘Base Excision Repair’ gene set (NES=2.75, q-value<0.001) and the Reactome ‘Double Strand Break Response’ (NES=2.05, q-value=0.0026). **(b)** Comparison of drug sensitivity (determined by IC50 values) to 72hr drug treatment with topotecan, between pseudo-diploid clones (SS48 and SS31) and highly-aneuploid clones (SS51 and SS111). n=5 (SS31) and n=6 (SS48, SS51, SS111) independent experiments. Fold change of IC50 calculated per experiment, relative to SS48. IC50 fold-change was calculated relative to SS48, per experiment. **, p=0.0099 (SS111); One-Sample t-test. **(c)** Gene set enrichment analysis (GSEA) of DNA damage response (DDR) gene expression signatures, comparing the highly-aneuploid clones, SS51 and SS11, to the pseudo-diploid clone SS48. Shown is an enrichment plot for the transcriptional targets of p53 (‘Kannan_TP53_Targets_Up’ gene set; NES=2.09, q-value=0.004). **(d)** Comparison of the mRNA expression levels of the p53 transcriptional targets, quantified by qRT-PCR, between pseudo-diploid clones (SS48 and SS31) and highly-aneuploid clones (SS51 and SS111): CDKN1A (p21), MDM2, TIGAR and RRM2B. n=5 independent experiments. Expression fold-change was calculated relative to SS48, per experiment. CDKN1A: *, p=0.0207, p=0.0104 and p=0.0282 for SS31, SS51 and SS111, respectively; MDM2: *, p=0.0175 and p=0.0315 for SS51 and SS111 respectively; TIGAR: *, p=0.0386 and **, p=0.0028 and p=0.0049 for SS31, SS51 and SS111 respectively; RRM2B: **, p=0.0024 (SS111); One-Sample t-test. **(e)** Immunofluorescence of _Y_H2AX foci in pseudo-diploid RPE1 cells, and their highly-aneuploid derivatives RPTs. Green, _Y_H2AX; Blue, DAPI; Scale bar, 20μm. **(f)** Quantitative comparison of _Y_H2AX foci between pseudo-diploid RPE1* and highly aneuploid RPT cells. n=6 independent experiments; *, p=0.0273 (RPT4/RPE1) and ****, p<0.0001 (RPT1/RPE1, RPT3/RPE1); One-Way ANOVA, Dunnett’s multiple comparison. **(g)** Comparison of drug sensitivity (determined by IC50 values) to 72hr drug treatment with etoposide, between pseudo-diploid RPE1 cells, and their highly-aneuploid derivatives RPTs. n=6 independent experiments. IC50 fold-change was calculated relative to RPE1, per experiment. *, p=0.0126, **, p=0.0072 and p=0.0095 for RPT1, RPT3 and RPT4, respectively; One-Sample t-test. **(h)** Comparison of drug sensitivity (determined by IC50 values) to 72hr drug treatment with topotecan, between pseudo-diploid RPE1 cells, and their highly-aneuploid derivatives RPTs. n=5 independent experiments. IC50 fold-change was calculated relative to RPE1, per experiment. *, p=0.0208, p=0.0145, p=0.016 for RPT1, RPT3 and RPT4 respectively; One-Sample t-test. **(i-j)** Differential drug sensitivities between near-euploid and highly-aneuploid human cancer cell lines, based on the large-scale CTD^2^ drug screen (**i**) and PRISM screen (**j**) ^42,44^. Data are taken from Cohen-Sharir *et al*^7^. Direct DNA damage inducers (alkylating and intercalating agents, anti-topoisomerases, and PARP inhibitors) are highlighted in orange. Highly-aneuploid cell lines are more resistant to this class of drugs. (**k**) Pre-ranked GSEA of mRNA expression levels showing that high aneuploidy levels are associated with upregulation of the DNA damage response (DDR) in human primary tumors. Shown is the enrichment plot of Reactome ‘Base excision repair’ (NES=2.00; q-value=0.001) and ‘DNA double strand repair’ (NES=2.43, qvalue<0.001) gene sets. Data were obtained from the TCGA mRNA expression dataset^86^.

**Extended Data Figure 6: Characterization of CRAF activity and dependency in pseudo-diploid vs. highly-aneuploid RPE1 clones (related to Figure 4)**

**(a)** Comparison of drug sensitivity (determined by IC50 values) to 72hr drug treatment with the CRAF inhibitor PLX7904, between pseudo-diploid clones (SS48 and SS31) and highly-aneuploid clones (SS51 and SS111). n=4 independent experiments. IC50 fold-change was calculated relative to SS48, per experiment. ***, p=0.0008, **, p=0.0029, for SS51 and SS111, respectively; One-Sample t-test. **(b)** Comparison of the mRNA expression levels of BRAF, quantified by qRT-PCR, between pseudo-diploid clones (SS48 and SS31) and highly-aneuploid clones (SS51 and SS111). n=5 independent experiments. Expression fold-change was calculated relative to SS48, per experiment. **, p=0.0037, *, p=0.022 for SS51 and SS111, respectively; One-Sample t-test. **(c)** Western blot of BRAF protein levels in RPE1 clones. GAPDH was used as housekeeping control. **(d)** Quantification of BRAF protein levels between pseudo-diploid clones (SS48 and SS31) and highly-aneuploid clones (SS51 and SS111). n=5 independent experiments. **, p=0.0054 for SS51; One Sample t-test. **(e)** Western blot of total CRAF protein levels in RPE1 clones treated with siRNA against CRAF (or associated scrambled siRNA) for 72hrs. CRAF activation levels are associated with the degree of DNA damage in the cells. Tubulin was used as housekeeping controls. **(f-l)** Live-cell imaging quantification of multiple cell parameters following CRAF inhibition using 10µM 8-Br-cAMP for 72h. Presented representative images (**f**) of cells treated with the drug. Comparison of doubling time (**g**), dry mass doubling time (**h**), cell area (**i**), cell perimeter (**j**), cell track speed (**k**) and cell velocity (**l**), between the pseudo-diploid clones (SS48 and SS31) and highly-aneuploid clones (SS51 and SS111). n=7 independent experiments. Fold-change calculated relative to DMSO-treated cells; One-Way ANOVA, Tukey’s multiple comparison. * p<0.05 ; ** p<0.01 ; *** p<0.001. **(m-n)** Flow cytometry-based analysis of cell death following exposure to 10μM 8-Br-cAMP for 72hr. Cells were stained with SYTOX™ Green Treatment with etoposide (2.5μM for 72hr) was used as a positive control (**m**). About 1% of 8-Br-cAMP-treated cells were SYTOX™-positive in both pseudo-diploid and highly aneuploid cells (**n**). n=4 independent experiments; n.s., p>0.05; One-way ANOVA with Tukey’s multiple comparison.

**Extended Data Figure 7: Validation of increased CRAF activity and dependency across various model systems (related to Figure 4)**

**(a)** Western blot of pCRAF and total CRAF protein levels in pseudo-diploid clones (SS48 and SS31) pre-treated with reversine (500nM) or with control DMSO for 20hrs to induce aneuploidy, then harvested 72hrs post wash-out. Tubulin used as housekeeping control. **(b)** Quantification of CRAF activation based on the pCRAF/CRAF ratio in the reversine-treated pseudo-diploid clones (SS48/SS31), calculated relative to the DMSO control per experiment. n=6 independent experiments. *, p=0.0124 and p=0.0162 for reversine-treated SS48 and SS31 cells, respectively; One Sample t-test. **(c)** Western blot of pCRAF and total CRAF protein levels in inducible *TP53*-KD RPE1 parental cells, pre-treated with reversine (500nM) or with control DMSO for 20hrs to induce aneuploidy, then harvested 72hrs post wash-out. Vinculin and GAPDH were used as housekeeping controls. **(d)** Quantification of CRAF activation based on the pCRAF/CRAF ratio in the reversine-treated *TP53*-KD RPE1 parental cells, calculated relative to the DMSO control per experiment. n=5 independent experiments. *, p=0.043 and p=0.0227 for reversine-treated sh-CTL and sh-p53, respectively; One Sample t-test. **(e)** Western blot of pCRAF and total CRAF protein levels in inducible *TP53*-KO RPE1 parental cells, pre-treated with reversine (500nM) or with control DMSO for 20hrs to induce aneuploidy, then harvested 72hrs post wash-out. Vinculin was used as housekeeping control. **(f)** Quantification of CRAF activation based on the pCRAF/CRAF ratio in the reversine-treated *TP53*-KO RPE1 parental cells, calculated relative to the DMSO control per experiment. n=6 independent experiments. *, p=0.043 and p=0.0227 for reversine-treated sg-CTL and sg-p53 respectively; One Sample t-test. **(g)** Western blot of CRAF protein levels in reversine-treated parental RPE1 cells, treated with siRNA against CRAF (or control siRNA) for 72hrs. Tubulin was used as a housekeeping control. **(h)** Western blot of phospho-CRAF and total CRAF protein levels in RPE/RPT cells. Tubulin was used as housekeeping control. **(i-j)** Quantification of pCRAF protein levels (**i**) and CRAF activation (**j**) based on the pCRAF/CRAF ratio in RPE/RPT cells. Elevated pCRAF levels without increase of the pCRAF/CRAF ratio. n=4 independent experiments. pCRAF levels: p=0.059, *, p=0.0281, **, p=0.0087, for RPT1, RPT3, and RPT4 respectively; One Sample t-test. **(k)** Comparison of drug sensitivity (determined by IC50 values) to 72hr drug treatment with the CRAF inhibitor TAK632, between pseudo-diploid clones (SS48 and SS31) and highly-aneuploid clones (SS51 and SS111). n=5 independent experiments. IC50 fold-change was calculated relative to RPE1 per experiment. **, p=0.0032, p=0.002, p=0.0023, for RPT1, RPT3, and RPT4 respectively; One-Sample t-test. **(l-m)** Comparison of BRAF (**l**) and CRAF (**m**) protein expression, between the top and bottom aneuploidy quartiles of human cancer cell lines (n=168 and 166 cell lines, for BRAF and CRAF, respectively). Data were obtained from the DepMap proteomic 22Q1 release^58^. Two-tailed Mann-Whitney test. **(n)** Comparison of drug sensitivity (determined by IC50 values) to 72hr treatment with a sub-lethal dose (200nM) of the CRAF inhibitor TAK632, or with DMSO control, in highly-aneuploid RPE1 clones (SS51 and SS111). Sub-lethal dose of CRAF inhibition had no impact on cell viability. n=6 independent experiments. IC50 fold-change was calculated relative to the DMSO-treated cells, per experiment; One-Sample t-test.

**Extended Data Figure 8: Generation of inducible *TP53*-KD and *TP53*-KO RPE1 cells (related to Fig. 4 and Fig. 5)**

**(a)** Western Blot showing variation of p53 protein levels upon nutlin-3a stimulation in inducible *TP53*-KD and *TP53*-KO RPE1 cells. GAPDH was used as housekeeping control. **(b)** Quantification of relative p53 protein levels upon nutlin-3a stimulation in inducible *TP53*-KD and *TP53*-KO RPE1-hTERT cells relative to their related controls. n=4 independent experiments. **, p=0.006 and ***, p=0.0009 for *TP53*-KD and *TP53*-KO, respectively; two-tailed t-test. **(c-i)**: Downregulation of various p53 transcriptional targets (CDK1A (**c**), BAX (**d**), PUMA (**e**), MDM2 (**f**), TIGAR (**g**), GADD45A (**h**), RRM2B (**i**)) following nutlin-3a stimulation in inducible *TP53*-KD and *TP53*-KO RPE1 cells, relative to their related controls. *, p<0.05, **, p<0.01, ***, p<0.001, ****, p<0.0001 for each comparison; two-tailed t-test.

**Extended Data Figure 9: Increased sensitivity of aneuploid cells to MEK and ERK inhibition (related to Fig. 5)**

**(a)** Comparison of drug sensitivity (determined by IC50 values) to 72hr drug treatment with the MEK inhibitor selumetinib, between pseudo-diploid clones (SS48 and SS31) and highly-aneuploid clones (SS51 and SS111). IC50 fold-change was calculated relative to SS48, per experiment. n=4 (SS31) and n=6 (SS48, SS51, SS111) independent experiments; ***, p=0.0007 for SS51; One-Sample t-test. **(b)** Western blot of pMEK1/2 and total MEK1/2 protein levels in inducible *TP53*-KD RPE1 parental cells, pre-treated with reversine (500nM) or with control DMSO for 20hrs to induce aneuploidy, then harvested 72hrs post wash-out. Tubulin and Vinculin were used as housekeeping controls. **(c)** Quantification of MEK1/2 activation based on the pMEK/MEK ratio in the reversine-treated *TP53*-KD RPE1 parental cells, calculated relative to DMSO control per experiment. n=4 independent experiments. *, p=0.0174 and p=0.0424 for reversine-treated sh-CTL and sh-p53, respectively; One Sample t-test. **(d)** Western blot of pMEK1/2 and total MEK1/2 protein levels in inducible *TP53*-KO RPE1 parental cells, pre-treated with reversine (500nM) or with control DMSO for 20hrs to induce aneuploidy, then harvested 72hrs post wash-out. Vinculin was used as housekeeping control. **(e)** Quantification of MEK1/2 activation based on the pMEK/MEK ratio in the reversine-treated *TP53*-KO RPE1 parental cells, calculated relative to DMSO control, per experiment. n=5 independent experiments; **, p=0.0095 and *, p=0.014 for reversine-treated sg-CTL and sg-p53, respectively; One Sample t-test. **(f)** Western blot of pERK1/2 and total ERK1/2 protein levels in inducible *TP53*-KD RPE1 parental cells, pre-treated with reversine (500nM) or with control DMSO for 20hrs to induce aneuploidy, then harvested 72hrs post wash-out. Vinculin and Tubulin were used as housekeeping controls. **(g)** Quantification of ERK1/2 activation based on the pERK/ERK ratio in the reversine-treated *TP53*-KD RPE1 parental cells, calculated relative to the DMSO control, per experiment. n=5 independent experiments; **, p=0.0043 and *, p=0.0337, for reversine-treated sh-CTL and sh-p53, respectively; One Sample t-test. **(h)** Western blot of pERK1/2 and total ERK1/2 protein levels in inducible *TP53*-KO RPE1 parental cells, pre-treated with reversine (500nM) or with control DMSO for 20hrs to induce aneuploidy, then harvested 72hrs post wash-out. GAPDH was used as housekeeping control. **(i)** Quantification of ERK1/2 activation based on the pERK/ERK ratio in the reversine-treated *TP53*-KO RPE1 parental cells, calculated relative to the DMSO control per experiment. n=4 independent experiments. **, p=0.0087 and *, p=0.0136, for reversine-treated sg-CTL and sg-p53, respectively; One Sample t-test. **(j)** Comparison of drug sensitivity (determined by AUC) to the MEK inhibitor selumetinib, between the top and bottom aneuploidy quartiles of human cancer cell lines (n=422 cell lines). Data were obtained from GDSC1 drug screen, DepMap portal 22Q1 release. **, p=0.0028; Two-tailed Mann-Whitney test. **(k)** Comparison of viability following 72hrs treatment with a sub-lethal dose (0.45nM) of the MEK inhibitor trametinib or DMSO, in highly-aneuploid RPE1 clones (SS51 and SS111). Sub-lethal dose of MEK inhibition had no impact on cell viability. n=5 independent experiments. Fold change in viability was calculated relative to DMSO-treated cells, per experiment; One-Sample t-test. **(l)** Visualization of the synergic effect of combining a sub-lethal dose of MEK inhibitor trametinib (0.45nM) with Etoposide. D, DMSO; T, trametinib 0.45nM; E, etoposide 2.5μM; T+E, combination trametinib 0.45nM and etoposide 2.5μM.

## Author contributions

U.B.-D. and S.S. jointly conceived the study, directed and supervised it. J.Z. and M.R.I. jointly designed and performed most of the experiments. J.Z., M.R.I., U.B.-D. and S.S. analyzed the data with inputs from all co-authors. Y.E. and E.R. assisted with bioinformatic analyses. G.D.F., A.S.K., S. M, S. V., K.L., Y. C.-S. and S.S. assisted with *in vitro* experiments. G.L. generated aneuploidy scores. J.B. performed the primary drug screen. F.V. directed the genomic profiling and CRISPR screens. J.Z., M.R.I., U.B.-D. and S.S. wrote the manuscript with inputs from all co-authors.

## Materials availability

Aneuploid RPE1-hTERT clones generated in this study are available upon request to Stefano Santaguida. Low-pass whole-genome sequencing and raw RNAseq data are available in the SRA database (https://www.ncbi.nlm.nih.gov/sra) under accession numbers PRJNA672256 and PRJNA889550, respectively. Whole-exome sequencing data and genome-wide CRISPR/Cas9 screening data of RPE1-hTERT clones are available in **Supplementary Tables 2-3 and 6** and in the DepMap database 21Q3 release (https://figshare.com/articles/dataset/DepMap_21Q3_Public/15160110). Drug screening data are available in **Supplementary Table 7** and in the Drug Repurposing Hub (https://clue.io/repurposing). Cancer cell line expression, CRISPR/Cas9 and RNAi data are available in the DepMap database 22Q1 release (https://figshare.com/articles/dataset/DepMap_22Q1_Public/19139906). All of them are publicly available as of the date of publication. Code extension to the aneuploidy score of cancer cell lines presented in Cohen-Sharir *et al*^7^ can be downloaded from https://github.com/BenDavidLab/Aneuploidy_Score_Code_2021.

## Supplementary information

**Supplementary Table 1: Summary of the newly-derived RPE1-hTERT cell line library**

Low-pass whole-genome sequencing (lp-WGS) based karyotyping of the 199 single-cell derived clones that were propagated successfully to give rise to RPE1 clones. Trisomies are denoted with the value ‘1’, monosomies are denoted with the value ‘-1’. The clones that were selected for omics profiling, perturbation screens and mechanistic studies are highlighted in yellow.

**Supplementary Table 2: Mutation profiling of RPE1-hTERT clones**

**Supplementary Table 3: Gene copy number alteration profiling of RPE1-hTERT clones**

Log2CN values are provided for each clone.

**Supplementary Table 4: Differential gene expression analysis of highly-aneuploid vs. pseudo-diploid RPE1-hTERT clones**

DESeq2 differential gene expression analysis of highly-aneuploid (SS51/SS111) versus pseudo-diploid clone SS48.

**Supplementary Table 5: Differential gene expression analysis of aneuploid vs. pseudo-diploid RPE1-hTERT clones**

DESeq2 differential gene expression analysis of all aneuploid clones (SS6/SS119/SS51/SS111) versus pseudo-diploid clone SS48.

**Supplementary Table 6: CRISPR/Cas9 screen results of RPE1-hTERT clones**

CERES essentiality scores for each gene in each RPE1 clones. Values<-1 represent gene essentiality.

**Supplementary Table 7: Pharmacological screen results of RPE1-hTERT clones**

Relative viability (%), determined by normalizing signal to the signal window of (DMSO control - inhibitor control), of each RPE1 clone upon exposure to 5,336 compounds. Shown are the average values of two technical duplicates. Compounds activity was defined by comparing the signal to 3SD of DMSO control. Compounds were defined as “active” if the mean of both replicates was equal or less than the activity threshold, as “inactive” if neither of the replicates was equal or less than the activity threshold, or “inconclusive” if one of the two replicates was equal or less than the activity threshold but the mean of both replicates was above the activity threshold. See **Methods** for more details.

**Supplementary Table 8: Extended table of aneuploidy scores for human cancer cell lines** Aneuploidy scores (AS) for 1742 human cancer cell lines. AS were determined as the number of chromosome arms that were gained or lost in each cell lines.

**Supplementary Table 9: Genes associated with high proliferation in highly-aneuploid, but not near-euploid, human cancer cell lines**

List of genes whose overexpression is significantly associated with high proliferation (i.e., low doubling time) in highly-aneuploid (AS≥21) cancer cell lines, but not in near-euploid (AS≤8) cancer cell lines.

**Supplementary Table 10: PRISM screen results of human cancer cell lines treated with selumetinib, in the absence or presence of reversine**

Comparison of the EC50 values of selumetinib in human cancer cell lines in the absence or presence of a sub-lethal dose (250nM) of reversine (or vehicle control) for 5 days.

**Supplementary Table 11: Details of reagents and oligonucleotides used in the study**

