## Supplementary figures and images for "Human aneuploid cells depend on the RAF/MEK/ERK pathway for overcoming increased DNA damage"

### Extended Data Fig. 1

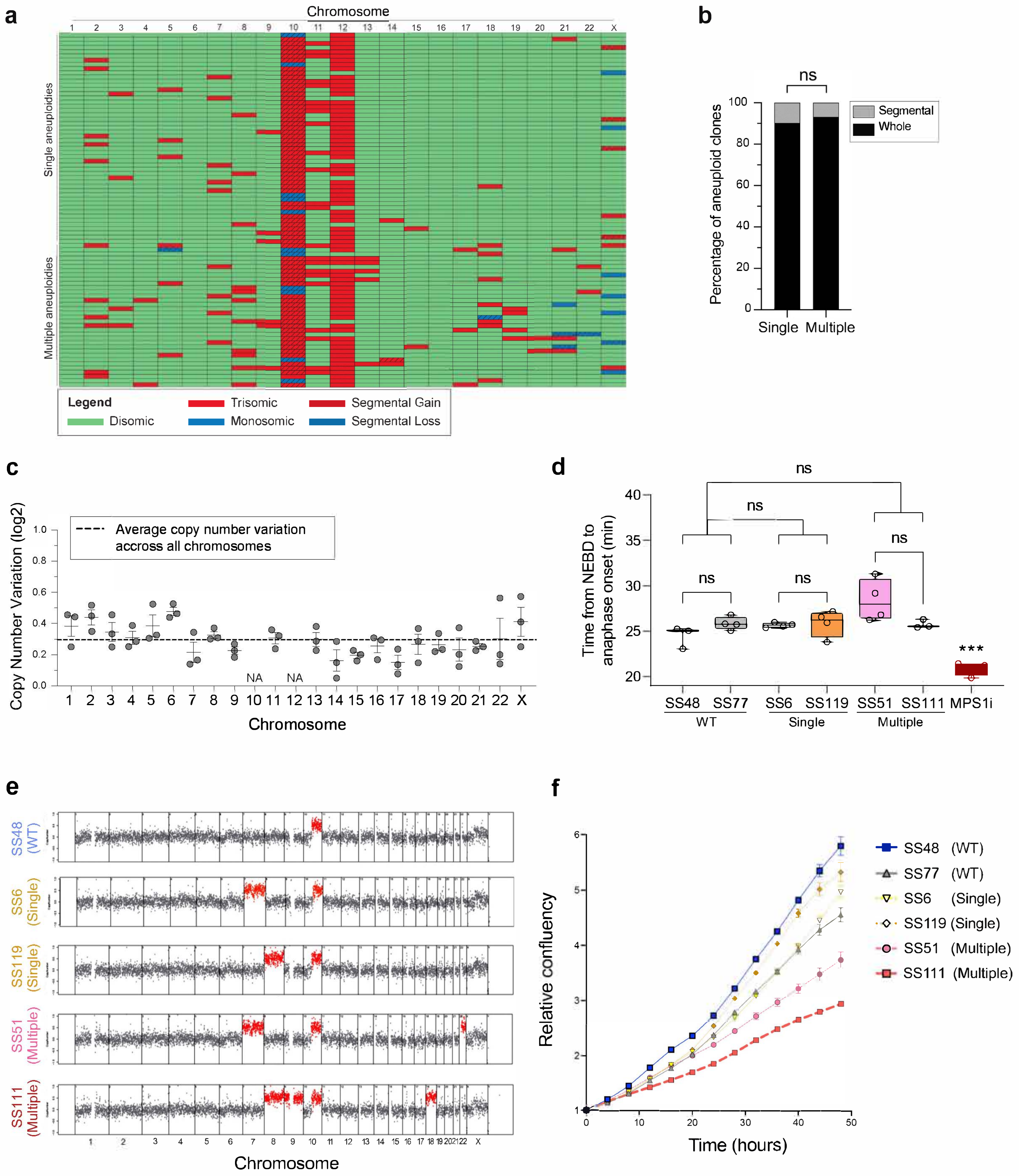


**Extended Data Figure 1**

### Extended Data Fig. 2

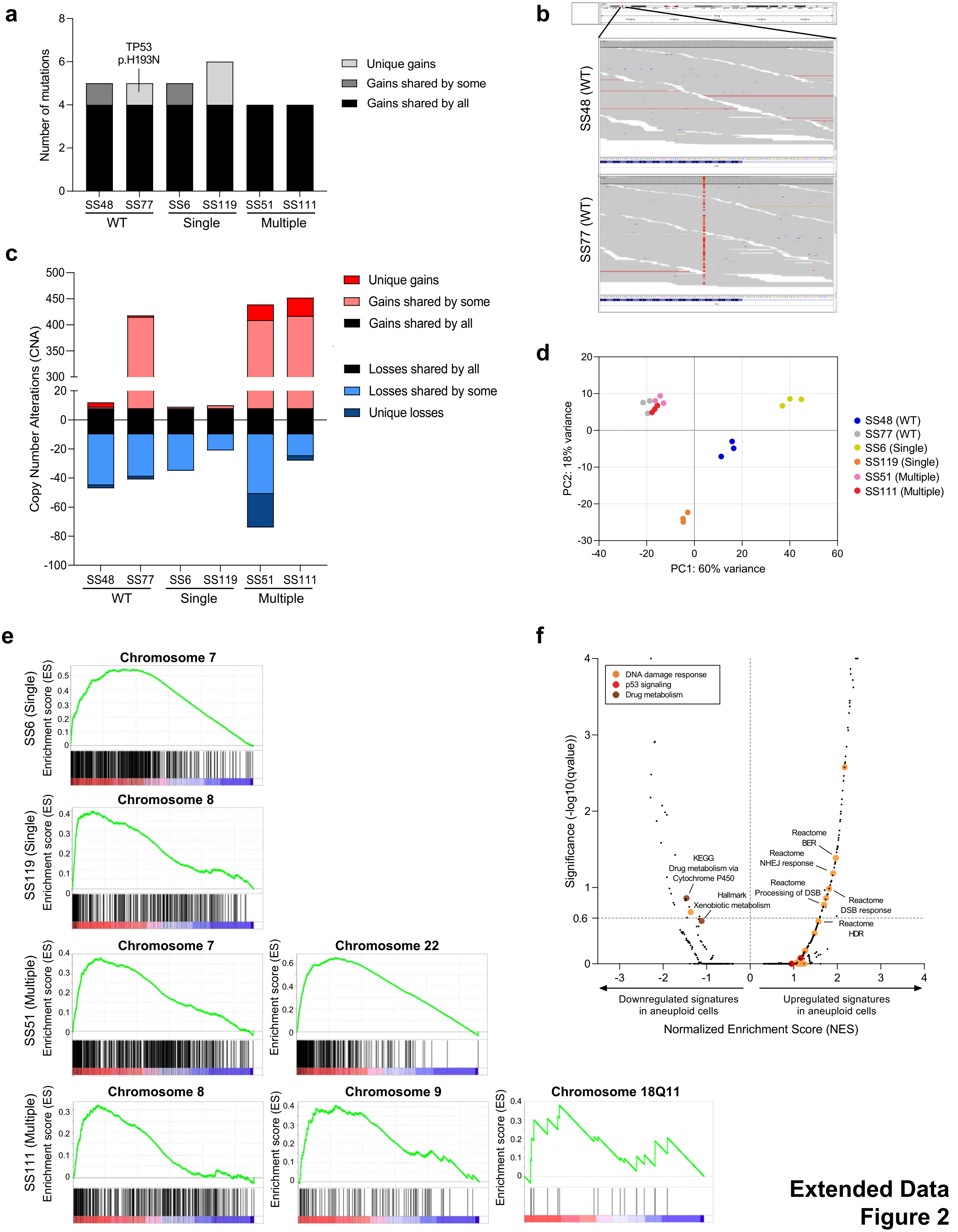


**Extended Data Figure 2**

### Extended Data Fig. 3

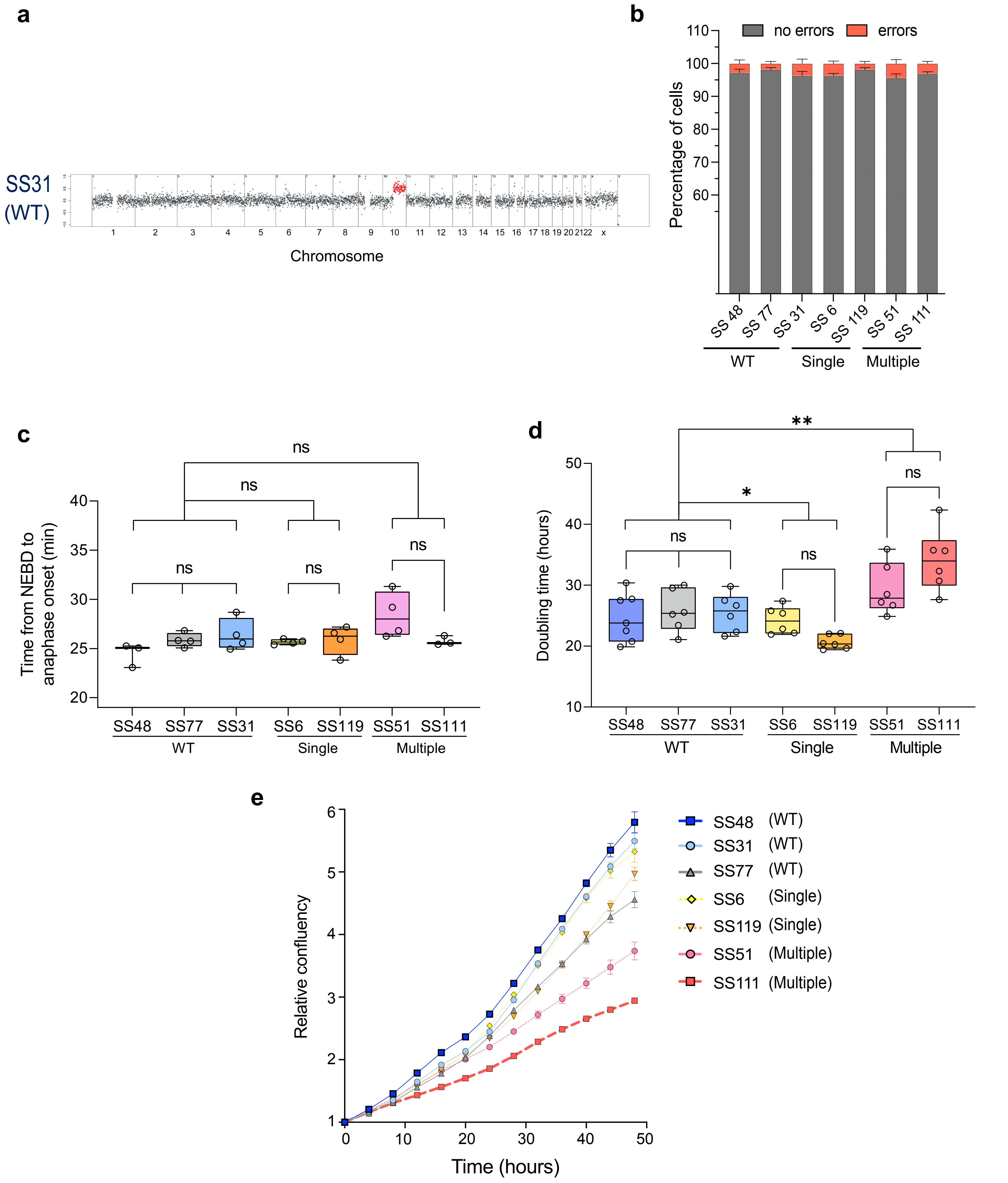


**Extended Data Figure 3**

### Extended Data Fig. 4

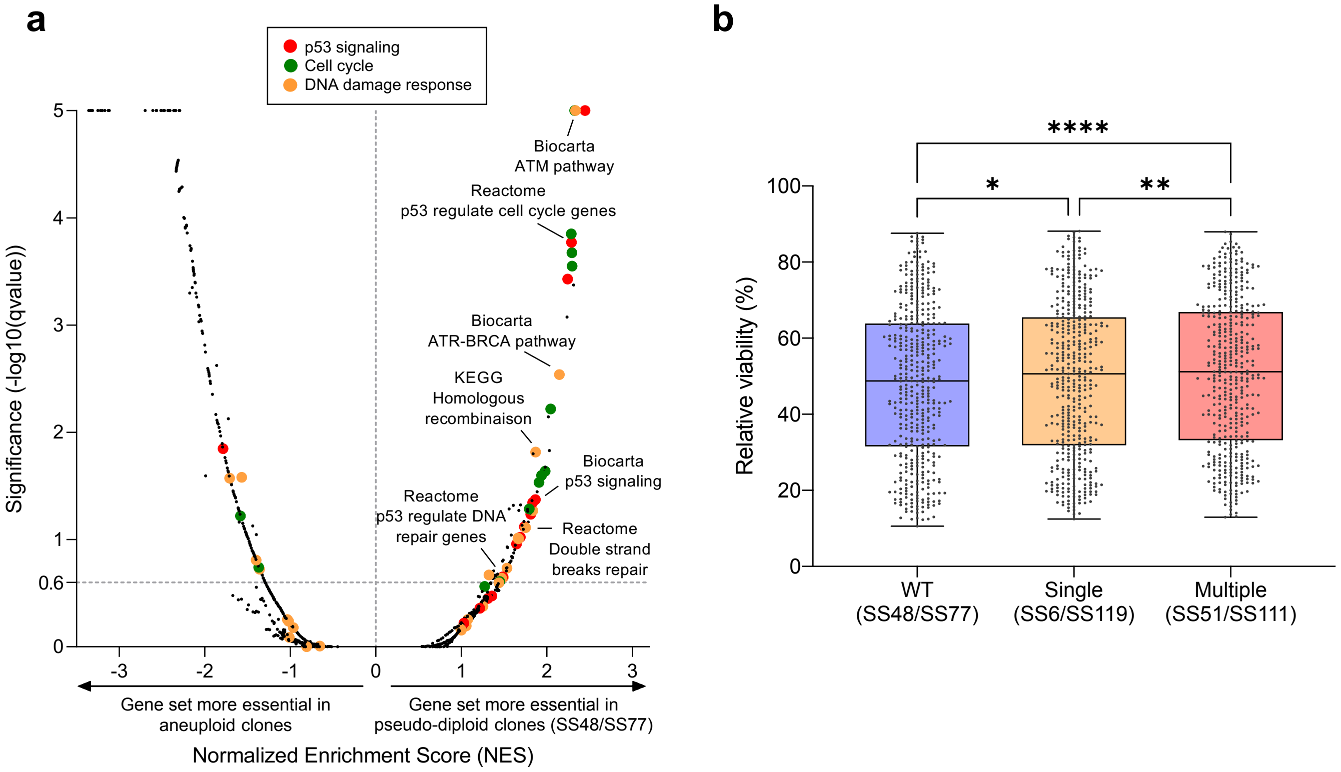


**Extended Data Figure 4**

### Extended Data Fig. 5

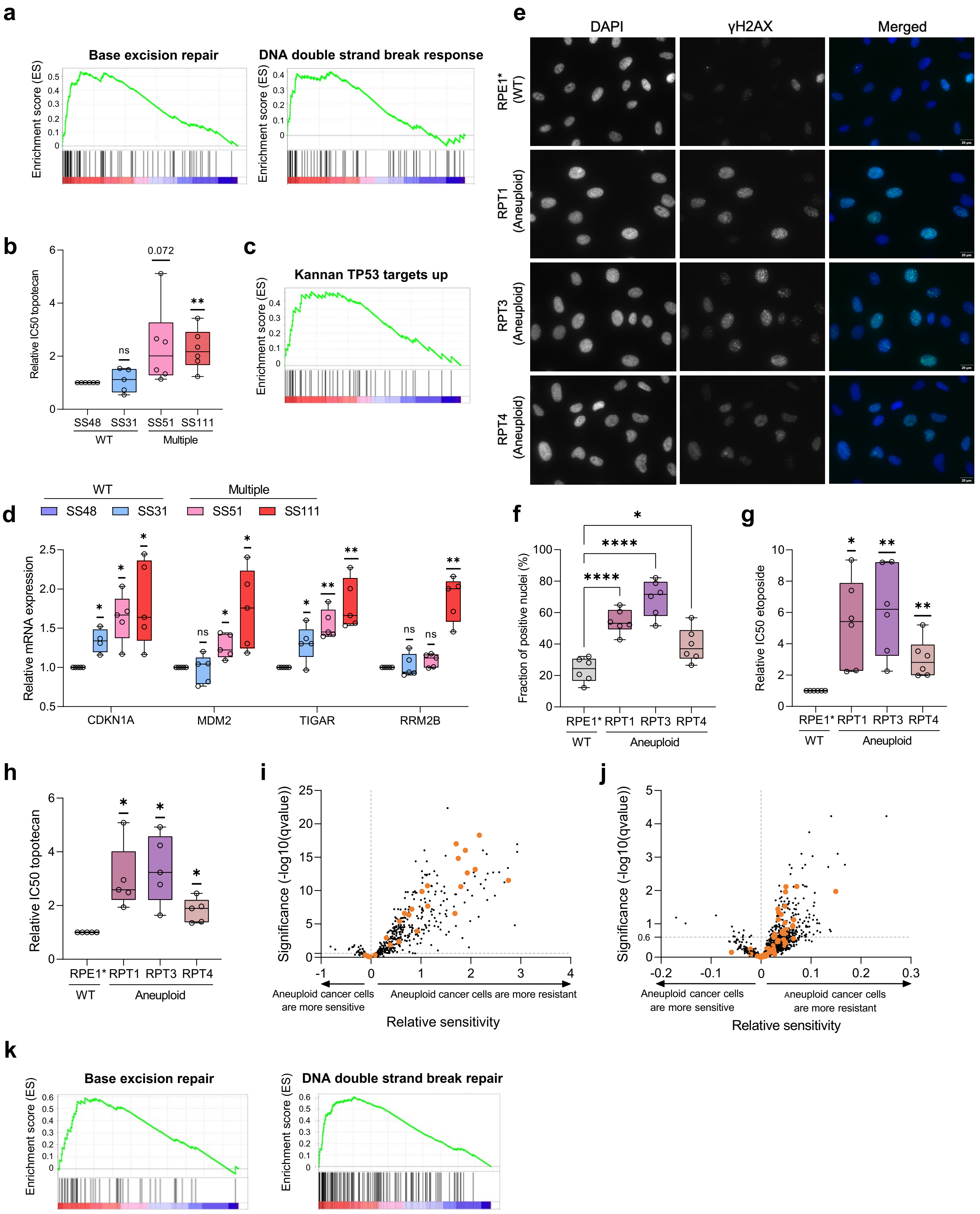


#### Extended Data Figure 5

### Extended Data Fig. 6

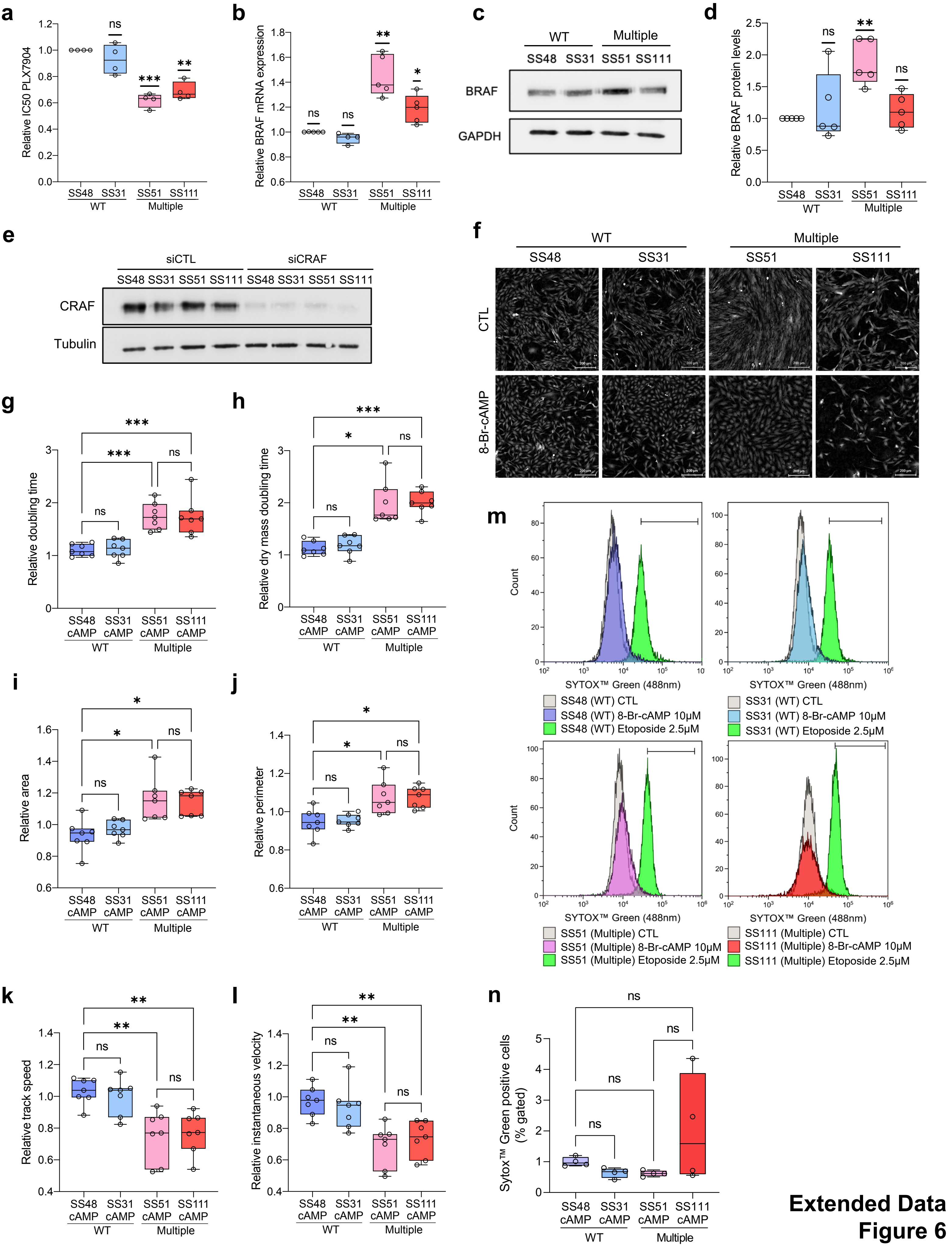


**Extended Data Figure 6**

### Extended Data Fig. 7

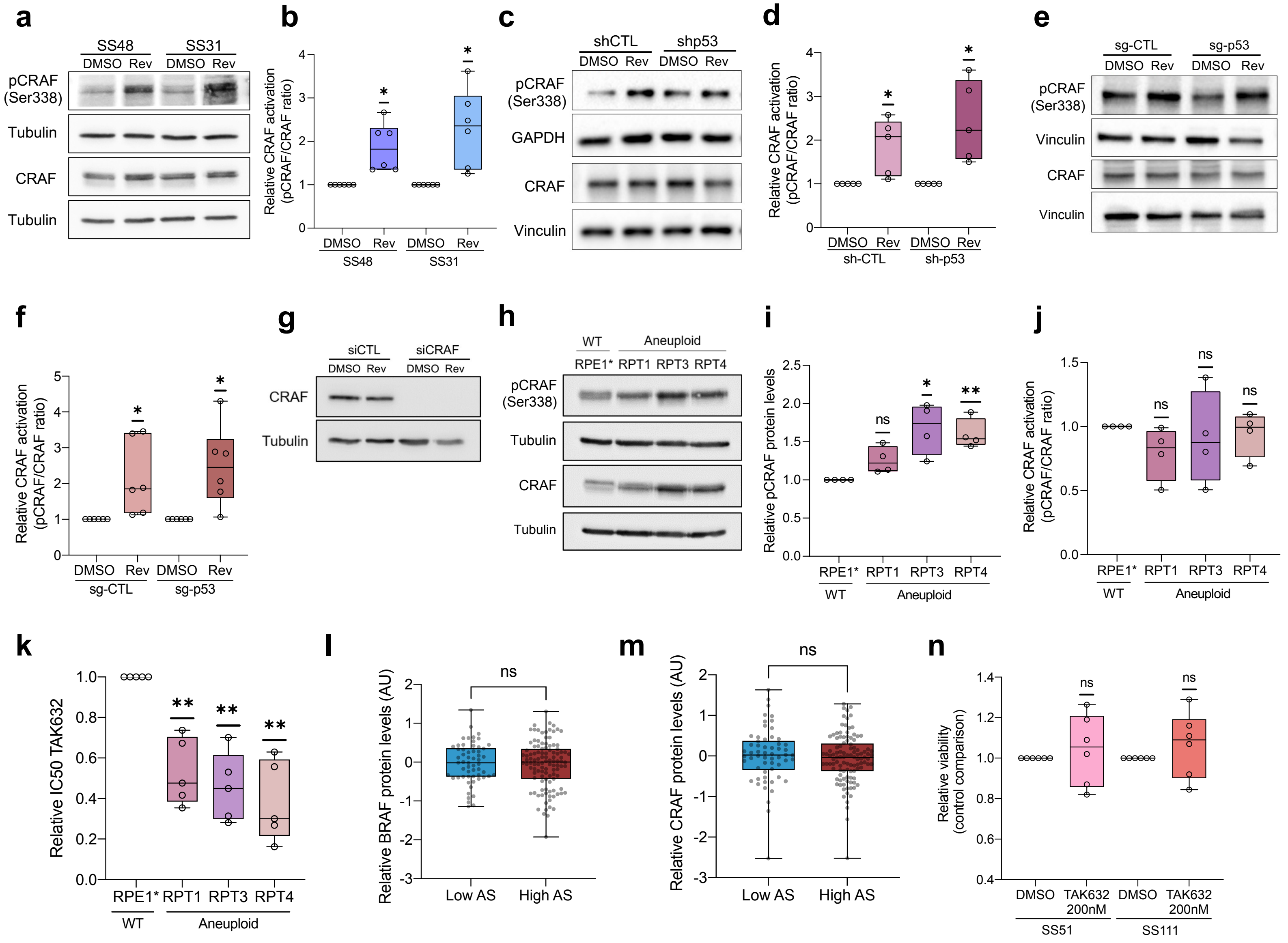


**Extended Data Figure 7**

### Extended Data Fig. 8

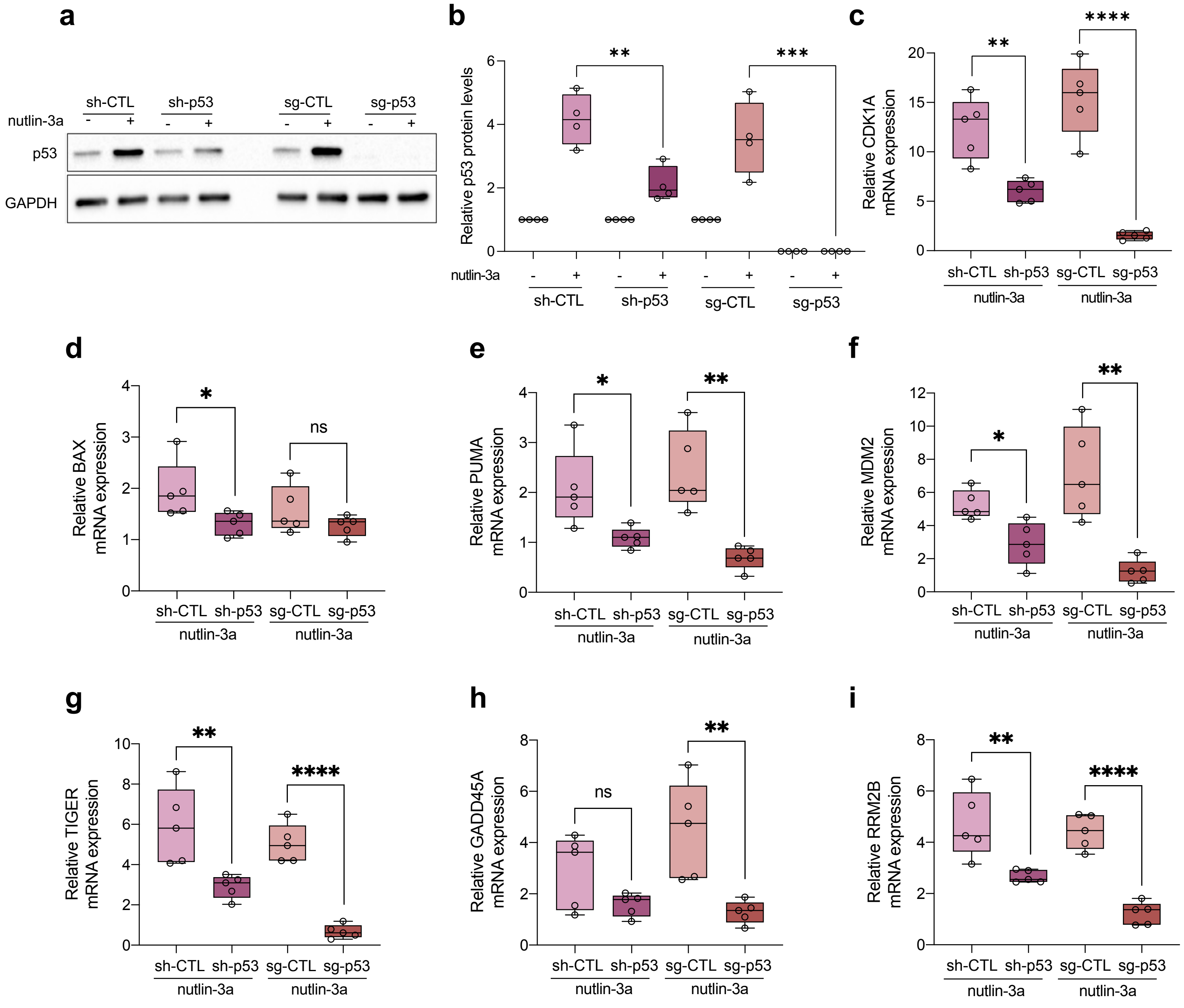


**Extended Data Figure 8**

### Extended Data Fig. 9

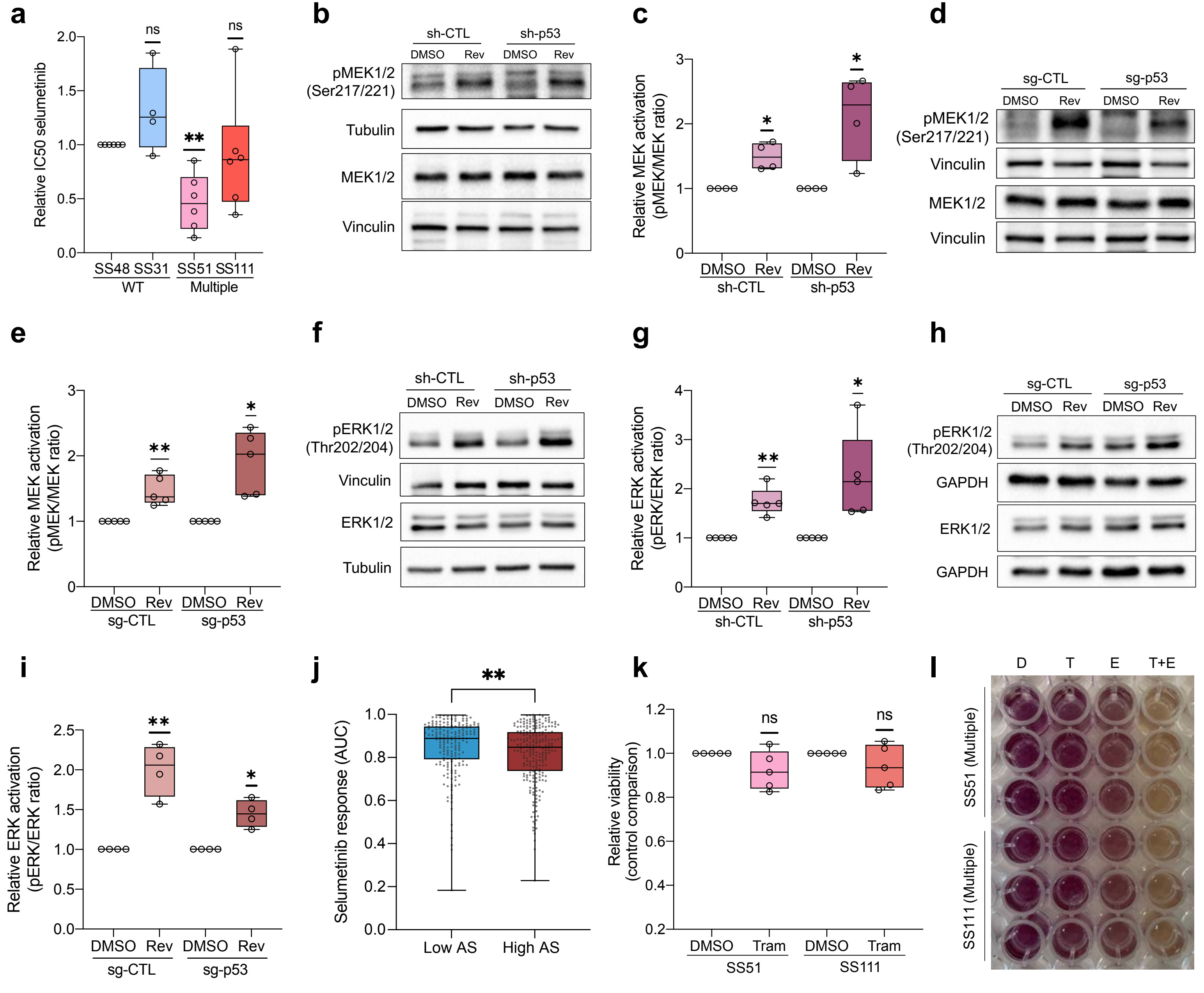


**Extended Data Figure 9**
